# Combined SHP2 and ERK inhibition for the treatment of KRAS-driven Pancreatic Ductal Adenocarcinoma

**DOI:** 10.1101/2021.12.14.472574

**Authors:** Katrin J. Ciecielski, Antonio Mulero-Sánchez, Alexandra Berninger, Laura Ruiz Cañas, Astrid Bosma, Kıvanç Görgülü, Nan Wu, Kalliope N. Diakopoulos, Ezgi Kaya-Aksoy, Dietrich A. Ruess, Derya Kabacaoğlu, Fränze Schmidt, Larissa Heinemann, Yuhui Fan, Bram Thijssen, Marieke van de Ven, Natalie Proost, Susanne Kossatz, Wolfgang A. Weber, Bruno Sainz, Rene Bernards, Hana Algül, Marina Lesina, Sara Mainardi

## Abstract

Mutant KRAS is present in over 90% of pancreatic as well as 30-40% of lung and colorectal cancers and is one of the most common oncogenic drivers. Despite decades of research and the recent emergence of isoform-specific KRAS^G12C^-inhibitors, most mutant KRAS isoforms, including the ones frequently associated with pancreatic ductal adenocarcinoma (PDAC), cannot be targeted directly. Moreover, targeting single RAS downstream effectors induces adaptive mechanisms leading to tumor recurrence or resistance. We report here on the combined inhibition of SHP2, a non-receptor tyrosine phosphatase upstream of KRAS, and ERK, a serine/threonine kinase and a key molecule downstream of KRAS in PDAC. This combination shows synergistic anticancer activity *in vitro*, superior disruption of the MAPK pathway, and significantly increased apoptosis induction compared to single-agent treatments. *In vivo*, we demonstrate good tolerability and efficacy of the combination. Concurrent inhibition of SHP2 and ERK induces significant tumor regression in multiple PDAC mouse models. Finally, we show evidence that ^18^F-FDG PET scans can be used to detect and predict early drug responses in animal models. Based on these compelling results, we will investigate this drug combination in a clinical trial (SHERPA, **SH**P2 and **ER**K inhibition in **pa**ncreatic cancer, NCT04916236), enrolling patients with *KRAS*-mutant PDAC.

## Introduction

Pancreatic cancer is the third leading cause of cancer-related deaths in western countries, the seventh world-wide, and is predicted to become the second most common cause of cancer mortality in the US in the next twenty to thirty years (1–3). The 5-year survival rate of patients suffering from PDAC is only 10% as diagnosis is often made when disease is already advanced, and therapeutic options are limited. Over the last years, the genomic landscape of PDAC has emerged, and recurrent mutations have been identified (4, 5). Recently, the discovery of a subset of PDAC patients bearing germline alterations in *BRCA1/2* and *PALB2*, which cause homologous repair deficiency (HRD), has led to the approval of PARP inhibitors as the first targeted therapy for HRD-pancreatic cancer (6). However, *BRCA1/2* and *PALB2* mutants are present in only 5–9% of PDAC patients (7), compared to over 90% of patients that carry tumors bearing *KRAS* mutations, and to date, no other non-cytotoxic, targeted therapies have been approved.

It has been widely demonstrated that *KRAS* mutations constitute an early initiating event in the pancreatic tumorigenic process (8), and that pancreatic adenocarcinomas retain a high dependency on RAS signaling (9, 10), thus making KRAS the ideal therapeutic target for pancreatic cancer. However, for more than three decades, research on direct targeting of RAS has proven to be a very challenging task. Only recently, KRAS^G12C^-specific inhibitors have entered clinical development (11–13), and some, like sotorasib and adagrasib have shown initial clinical responses (14–16), leading to sotorasib being the first drug approved by the Food and Drug Administration (FDA) for the treatment of KRAS^G12C^ driven non-small cell lung cancer.

Unfortunately, the G12C variant represents only 1% of *KRAS* mutations in pancreatic cancer, with the most frequent amino acid substitutions being G12D (41%), G12V (34%) and G12R (16%) (17). For those most common mutations, targeted inhibitors have not yet been developed; therefore, most translational studies have been aimed at blocking downstream RAS effectors mainly in the MAPK or PI3K-AKT pathways (18–21). Unfortunately, attempts to target RAS downstream effectors have been hampered by compensatory feedback mechanisms, often involving reactivation of receptor tyrosine kinases (22). This notion, together with advances in KRAS biophysics and structural biology studies (23, 24) undermined the old paradigm of mutant KRAS being constitutively active and made it clear that it is possible to reduce mutant RAS activation by combining inhibition of upstream and downstream nodes in the RAS-MAPK pathway. In particular, the ubiquitously expressed non-receptor protein tyrosine phosphatase SHP2, encoded by the *PTPN11* gene, has been identified as a useful upstream target (25–28), as it is involved in signal transduction downstream of multiple growth factor, cytokine, and integrin receptors (29). Potent and specific allosteric inhibitors of SHP2 have recently been developed and have entered clinical trials (30), holding great promise for receptor tyrosine kinase (RTK)-driven tumors. Nonetheless, so far, the question regarding the most beneficial drug combination for KRAS-driven pancreatic cancer remains.

MEK inhibitors have been tested extensively in *KRAS*-mutant PDAC as well as other solid tumors with poor results (31–33). This is primarily attributable to their highly toxic profile and adverse side effects as well as the above-mentioned feedback reactivation of the MAPK pathway (22, 34). Inhibitors of ERK, directly downstream of MEK, have only recently been introduced into clinical studies (35) and seem auspicious with regard to their toxicity profile. In the present study, we explore the tolerability and efficacy of combining the allosteric SHP2 inhibitor RMC-4550 with the ATP-competitive, selective ERK inhibitor LY3214996 in multiple *in vitro* and *in vivo* models of murine and human PDAC. Based on the data reported here, we developed the Phase 1a/1b SHERPA clinical trial (NCT04916236).

## Results

### Combinatorial effect *in vitro*

In a previous study, we showed promising *in vitro* and *in vivo* results to support combined SHP2 and MEK inhibition for the treatment of pancreatic cancer (25, 27). While the reported combinatorial strategy is sound, the frequently observed side effects and the common occurrence of resistance associated with MEK inhibitors like trametinib (34, 36, 37), led us to search for possible alternatives. Since reactivation of ERK is a major mechanism hampering MEK inhibitor (MEKi) efficacy (22, 36, 38, 39), we hypothesized that direct ERK inhibition might be a worthwhile strategy. The newly developed ERK inhibitor LY3214996 (40) was shown to be a selective, potent, and reversible ATP-competitive inhibitor of ERK1/2 activity in KRAS- and BRAF-mutant cell lines. In parallel, a selective allosteric SHP2 inhibitor (RMC-4550) was developed with a mode of action similar to the Novartis’ SHP099, but with slightly higher potency (26).

Since LY3214996 (ERKi) and RMC-4550 (SHP2i) have not yet been studied in combination, we first investigated the effects of combined treatment on MAPK activity, proliferation, and apoptosis *in vitro*. First, we analyzed the inhibitors’ capacity to inhibit MAPK pathway activity. MEKi monotherapy is known to induce only a transient suppression of downstream MAPK signaling, before feedback RTK reactivation restores it to the initial levels (25), which can be prevented by concomitant SHP2 inhibition. To test whether a similar MAPK signal dynamic is induced by ERK inhibition, murine and human *KRAS*-mutant PDAC cell lines were treated with either ERKi alone or ERKi + SHP2i for 6, 72, and 144 hours. Western blot analysis was performed to examine protein expression levels of phosphorylated Ribosomal S6 kinase 1 (pRSK-1), a direct downstream target of ERK (41). Compared to ERKi monotherapy, combined ERKi + SHP2i treatment inhibited MAPK pathway activity more effectively at all time points analyzed (**Figure 1 A**). After 6 days (144 h), a 97%, 81%, and 81% reduction in pRSK-1 levels in comparison to the control could still be observed for ERKi + SHP2i-treated murine *Kras;Trp53*^-/-^ (KCP) K2101 as well as human MiaPaCa-2, and Panc10.05 cells, respectively, compared to only 21%, 43%, and 59% reduction in the same cells with ERKi monotherapy.

**Figure 1.**
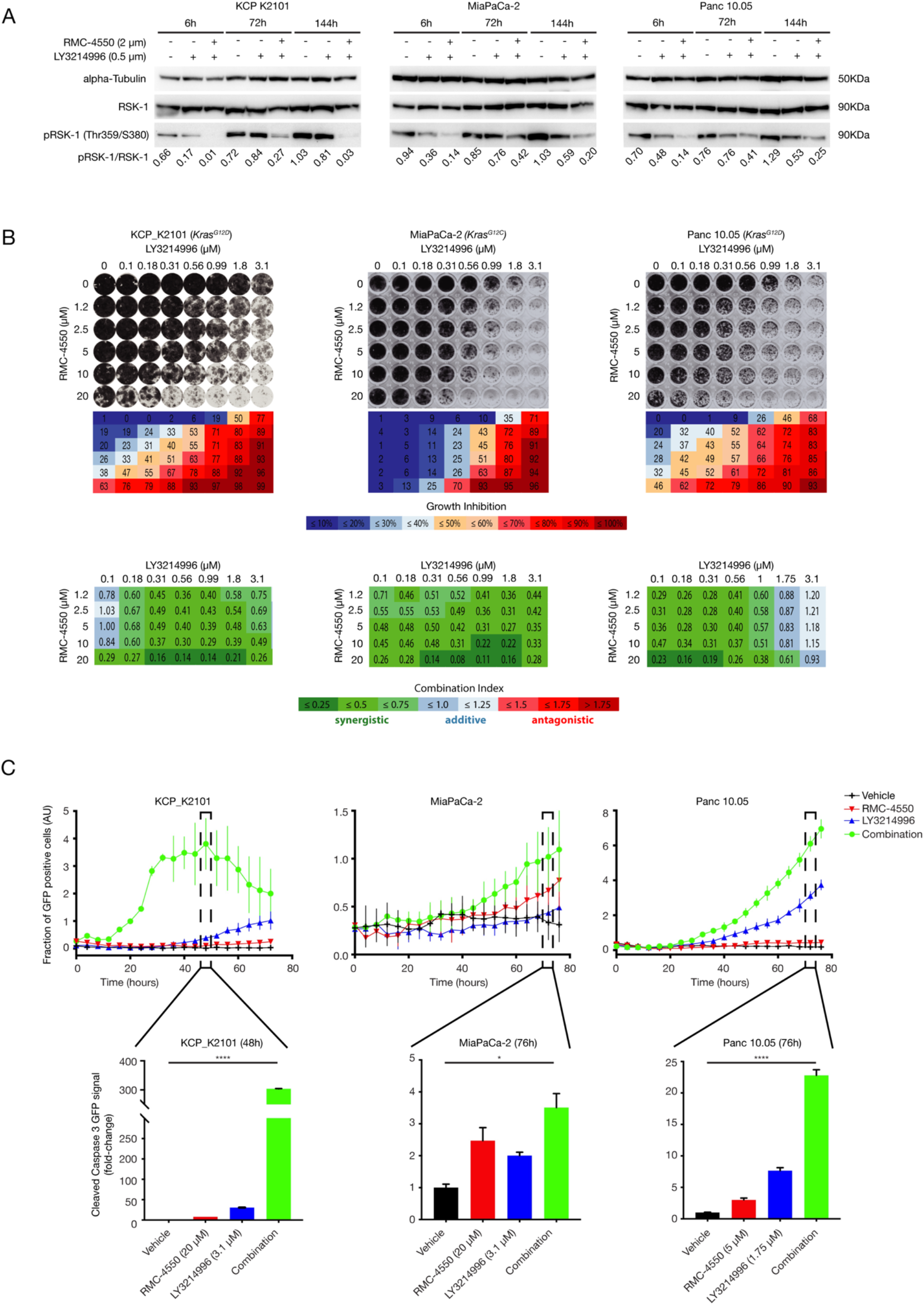
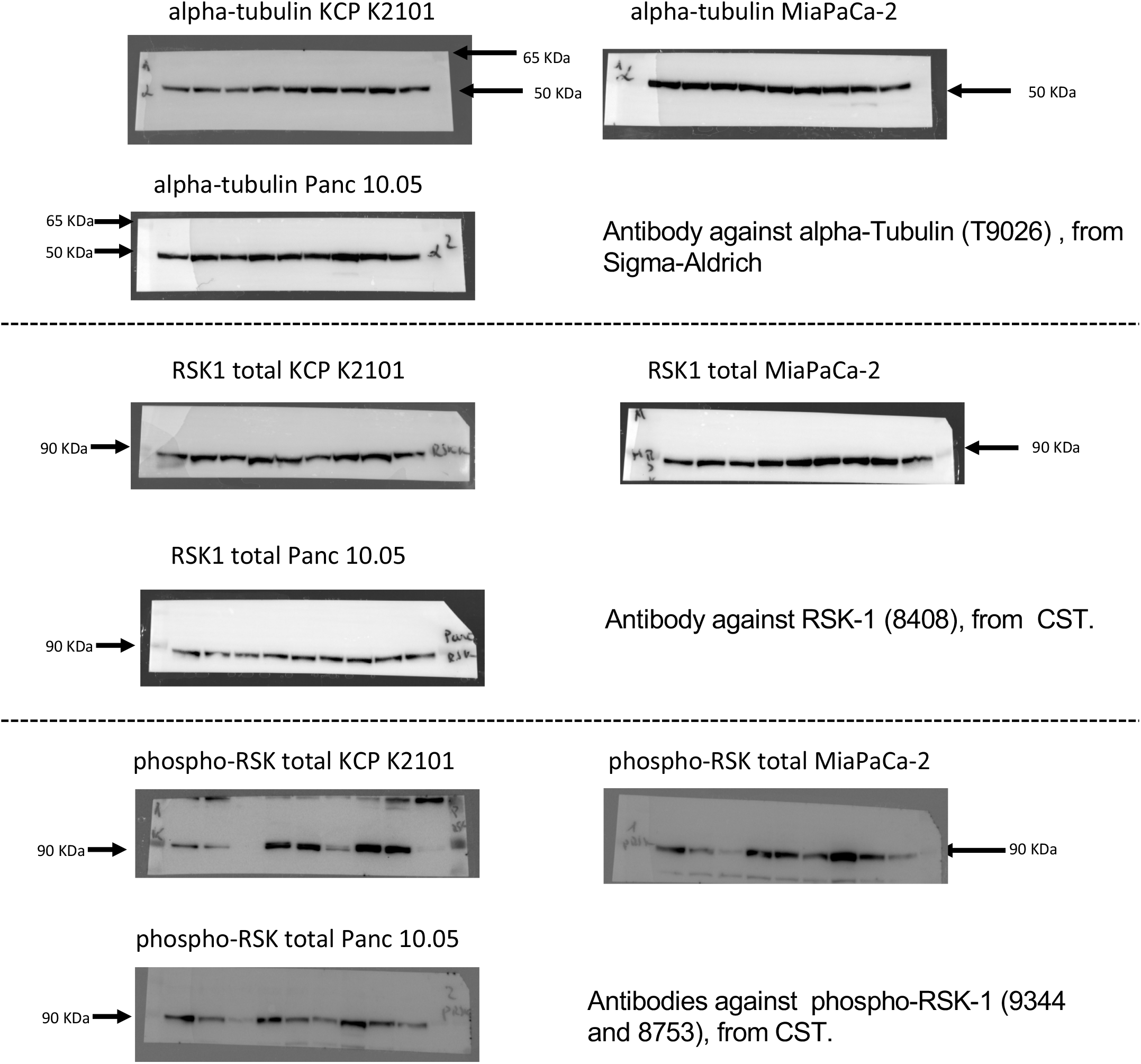
Evaluation of the combined effects of RMC-4550 (SHP2i) and LY3214996 (ERKi) administration in murine and human *KRAS*-mutant pancreatic cancer cell lines. **A:** Western blot analysis with murine cancer cell line KPC_K2101 derived from KPC mouse model (KRAS^G12D^) of spontaneous tumor formation and in human cancer cell lines: MiaPaCa-2 (KRAS^G12C^) and Panc 10.05 (KRAS^G12D^). Cells were treated as depicted and collected for lysis at the indicated time points. Protein extracts were probed with specific antibodies against total RSK-1, phosphorylated RSK-1 (pRSK-1), and alpha-tubulin (as loading control). Numerical values indicate the pRSK-1/RSK-1 ratio quantified by densitometry. The blots are representative of at least three independent experiments. RSK-1 = Ribosomal s6 kinase 1. **B:** Synergistic effects of SHP2i and ERKi administration were evaluated by colony formation assay in the *KRAS*-mutant cell lines used in **A**. SHP2i and ERKi were combined at the indicated concentrations. Representative crystal violet staining of cells is shown (top panel). Box-matrices below the plate-scans depict quantification of growth inhibition in relation to control wells (middle panel). Bottom panel: Calculation of the Combination Index (CI) Scores from the growth inhibition values (shown above) via CompuSyn software demonstrating strong synergism between SHP2i and ERKi across a wide range of combinatorial concentrations. CI < 0.75 (shades of green) indicates synergism, CI = 0.75 - 1.25 (shades of blue) indicates additive effects and CI > 1.25 (shades of red) indicates antagonism. Experiments were repeated independently at least three times each, with similar results. **C:** Apoptosis was analyzed in cell lines treated with either DMSO, SHP2i alone, ERKi alone, or combination of SHP2i and ERKi at the indicated concentrations in real time (top panel). GFP signal coupled to cleaved caspase 3 was quantified as read-out. Bar plots for selected time points (48 hours for KPC_K2101 and 76 hours for MiaPaCa-2 and Panc 10.05) show the fraction of GFP positive cells (AU) (top panel) and the fold-change GFP signal (bottom panel). AU = Arbitrary Units, GFP = Green Fluorescent Protein. Experiments were repeated independently at least three times each. Results represent mean ± SD. * P < 0.05, **** P < 0.0001, as determined by ordinary one-way ANOVA test.

To better understand the potential clinical benefit of the combination therapy, we further tested the effect of ERKi + SHP2i treatment on cell proliferation (**Figure 1 B and Supplementary Figure S1**) and on the induction of cell death. For the former, we performed 6-day colony formation assays using two murine KCP and 5 different human PDAC cell lines, harboring KRAS^G12C^, KRAS^G12V^ or KRAS^G12D^ mutations (**Figure 1 B and supplementary Figure S1**). The 96-well format matrix allowed us to test a range of inhibitor concentrations both as a monotherapy as well as in combination. Synergism (i.e., a combination index (CI) score below 0.75, indicated in shades of green in **Figure 1 B and S1**) was observed in at least 70% of all inhibitor combinations tested in all 7 cell lines, regardless of the type of KRAS mutation present. These data indicate that LY3214996 and RMC-4550 can synergistically inhibit PDAC cell growth *in vitro* at micromolar concentrations.

To study the effect of the drug combination on apoptosis, KCP-K2101, MiaPaCa-2, and Panc10.05 cells were treated with either SHP2i or ERKi alone at roughly IC50 concentrations, or with the combination of the two inhibitors for up to 76 h, and the rate of apoptosis was measured by tracking GFP-labeled cleaved caspase 3 over time (**Figure 1 C, upper panels**). In murine KCP-K2101 cells, GFP positive cleaved caspase 3 levels peaked after 48 h, and the fraction of caspase 3 positive cells was significantly higher in the combination treated group compared to vehicle or to ERKi and SHP2i monotherapy. Similarly, human cell lines MiaPaCa-2 and Panc10.05 showed a peak of GFP-coupled to a cleaved caspase 3/7 specific recognition motif after 76 h, with a significant increase in combination-treated cells compared to either vehicle or monotherapies indicating that apoptosis was triggered significantly with the combination. Based on the kinetics of the different cell lines, we identified the time point where apoptosis was maximally triggered to calculate the fold-change in apoptosis for the monotherapies and combination treatment compared to vehicle (**Figure 1 C, lower panels**).

Taken together, our data show that LY3214996 and RMC-4550 act synergistically to both inhibit PDAC cell proliferation and MAPK signaling as well as induce significant levels of apoptosis *in vitro*, which prompted us to test the combination treatment *in vivo.*

### *In vivo* tolerability

Due to the novelty of both LY3214996 and RMC-4550, the scarce *in vivo* data available, as well as the non-existent data on combined toxicity, we performed a tolerability study in nontumor bearing wild-type (*Kras^LSLG12D^; Trp53^flox/flox^ - no Cre*) and NOD-scid gamma (NSG) mice to determine the maximum tolerated dose (MTD) of combined LY3214996 and RMC-4550. The inhibitors were administered once per day via oral gavage for 14 consecutive days (**Figure 2 A**). Following dosing recommendations by Eli Lilly and Revolution Medicines, we determined 9 different doses of inhibitor combinations labeled d1 (lowest) through d9 (highest), as illustrated in **Figure 2 B**. We applied a modified “3 + 3” study design (42, 43) using cohorts of three animals per dose. As shown in **Figure 2 C**, the first cohort was treated at a starting dose, and the subsequent cohorts were treated with ascending or descending doses according to the observed response. Dosing was increased until one or more mice per cohort experienced dose-limiting toxicities (DLT). In case two or more mice experienced dose-limiting toxicity, the dose escalation was stopped and the next lower dose, with no more than 1 in 6 mice showing signs of DLT, was determined as the MTD. If only one in three mice experienced DLT, the cohort was expanded to 6 mice and the dose-escalation continued if none of the additional three mice showed signs of DLT, otherwise the previous dose was determined as the MTD. Endpoints used as signs of DLT were weight loss of more than 20%, clinical score (abnormal behavior, signs of physical discomfort) and death. These parameters were evaluated daily, and animals were euthanized if either of these endpoints were met. Due to ethical and practical considerations, and to minimize the number of mice in the experiment, dose d5 was chosen as the starting dose. **Figure 2** shows the body weight profile of both wild-type (**Figure 2 D**) and NSG (**Figure 2 E**) mice over the course of 14 days of treatment. All doses were well tolerated in wild-type mice (**Figure 2 D**) while dose d9 (i.e., 100 mg/kg LY3214996 + 30 mg/kg RMC-4550) caused dose-limiting weight loss in NSG mice (**Figure 2 E**).

**Figure 2.**
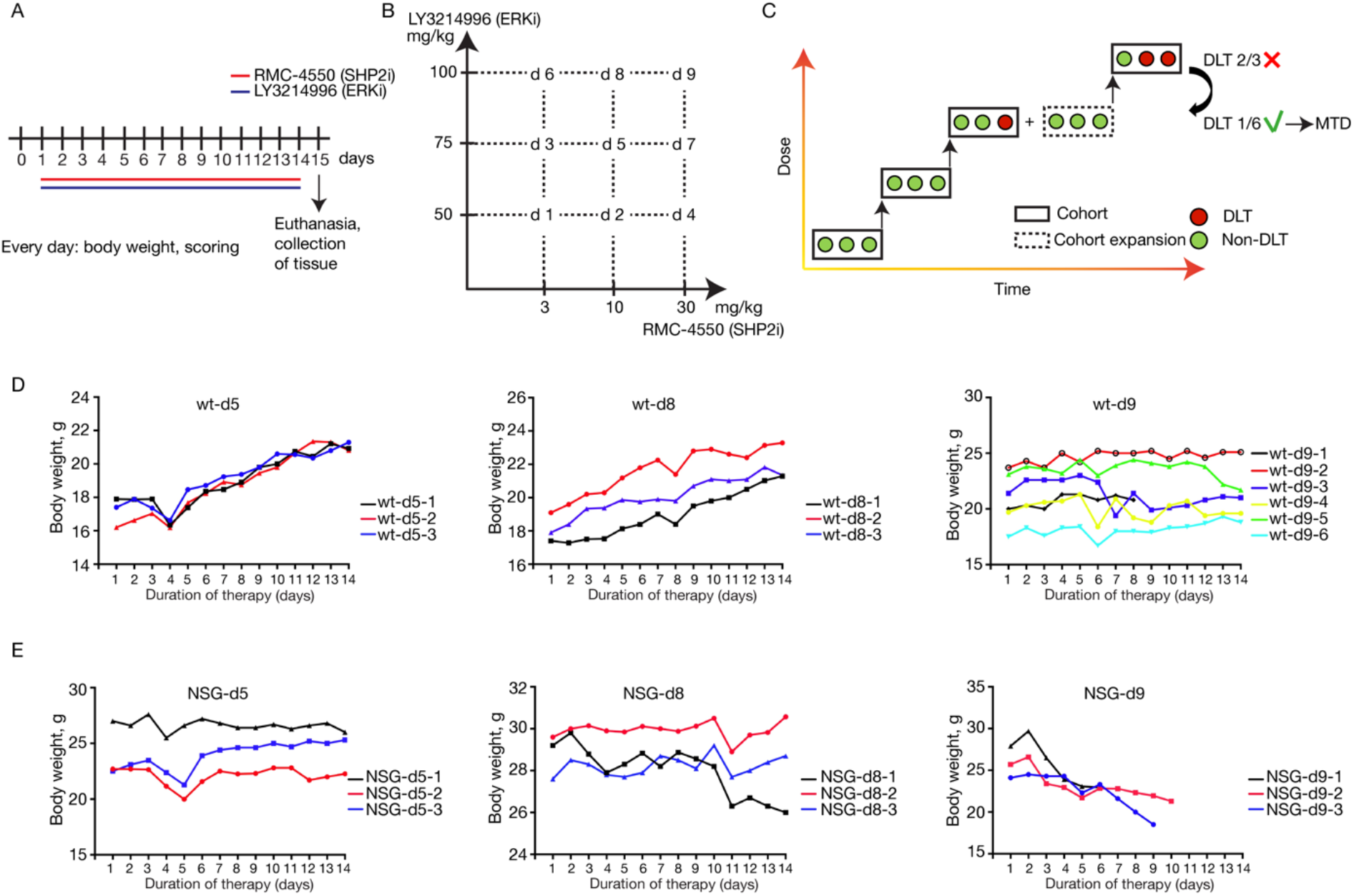
MTD Study design. **A**: Treatment schedule. Non-tumor bearing wild type and NOD scid gamma (NSG) mice were treated with the combination of RMC-4550 (SHP2i) and LY3214996 (ERKi) once per day via oral gavage for 14 consecutive days. **B**: Graphical representation of dose combinations. SHP2i and ERKi were combined in different concentrations to make up 9 combined doses. **C**: An illustration of the modified “3 + 3” study design. Each box represents a cohort comprising the indicated number of mice treated at a given dose level. DLT = Dose-Limiting Toxicity, MTD = Maximum Tolerated Dose. **D**: Individual body weight-time profile of the treatment groups in male wild type (wt) mice: d5 (n = 3), d8 (n = 3), d9 (n = 6). **E**: Individual body weight-time profile of the treatment groups in male NSG mice: d5 (n = 3), d8 (n = 3), d9 (n = 6).

Thus, dose d8, i.e., 100 mg/kg ERKi + 10 mg/kg SHP2i was the highest dose that was well tolerated in both wild-type and NSG mice and was therefore used to assess anti-tumor efficacy.

### *In vivo* efficacy

Having found a well-tolerated dose for the combination therapy of LY3214996 + RMC-4550, we investigated its potential anti-tumor efficacy *in vivo.* First, we used a xenograft model of subcutaneously transplanted human PDAC cell lines (**Figure 3 A and B**). Mice bearing tumors with a volume of approximately 200 mm^3^ were randomly assigned into either the vehicle, RMC-4550 (A), LY3214996 (B) or combination (C) cohort and were treated daily for 21 days via oral gavage (**Figure 3 A**). While RMC-4550 alone was already partially effective at reducing MiaPaCa-2 xenograft tumor growth compared to vehicle, tumor volume reduction of more than 30% in 12 out of 16 mice was only achieved upon continuous combination treatment. We were able to confirm these results in a model of orthotopically transplanted *Kras^G12D/+^; Trp53^R172H/+^; Pdx-1Cre* tumors in immunocompetent C57BL/6J mice (KCP^mut^). Specifically, following post-surgical tumor expansion over 2 weeks, mice were randomized into either the baseline, vehicle, SHP2i (A), ERKi (B) or combination (C) cohort. Baseline mice were sacrificed to confirm the presence of well-integrated and uniform orthotopic tumors. All other mice were treated for 14 days via oral gavage as shown in **Figure 3 A**. As **Figure 3 C** and **D** show, significant inhibition in tumor growth, seen macroscopically and indicated by decreased tumor weight, was observed in both monotherapy groups as well as in the combination treatment group compared to the vehicle cohort. No difference in treatment efficacy was observed between weigh-matched male and female mice (**Figure 3 D**). Notably, and in agreement with the synergy described *in vitro*, the combination therapy (Cohort C) was the most effective and induced a significantly stronger tumor volume reduction compared to ERKi or SHP2i monotherapies (**Figure 3 C-D**). To confirm the on-target activity of the combination therapy, we show substantial reduction in transcriptional-based MAPK pathway activity score (44) *in vivo* in both the orthotopic KCP^mut^ tumors as well as the subcutaneous MiaPaCa-2 xenografts (**Figure 3 E**) treated with continuous ERKi + SHP2i compared to vehicle-treated controls.

**Figure 3.**
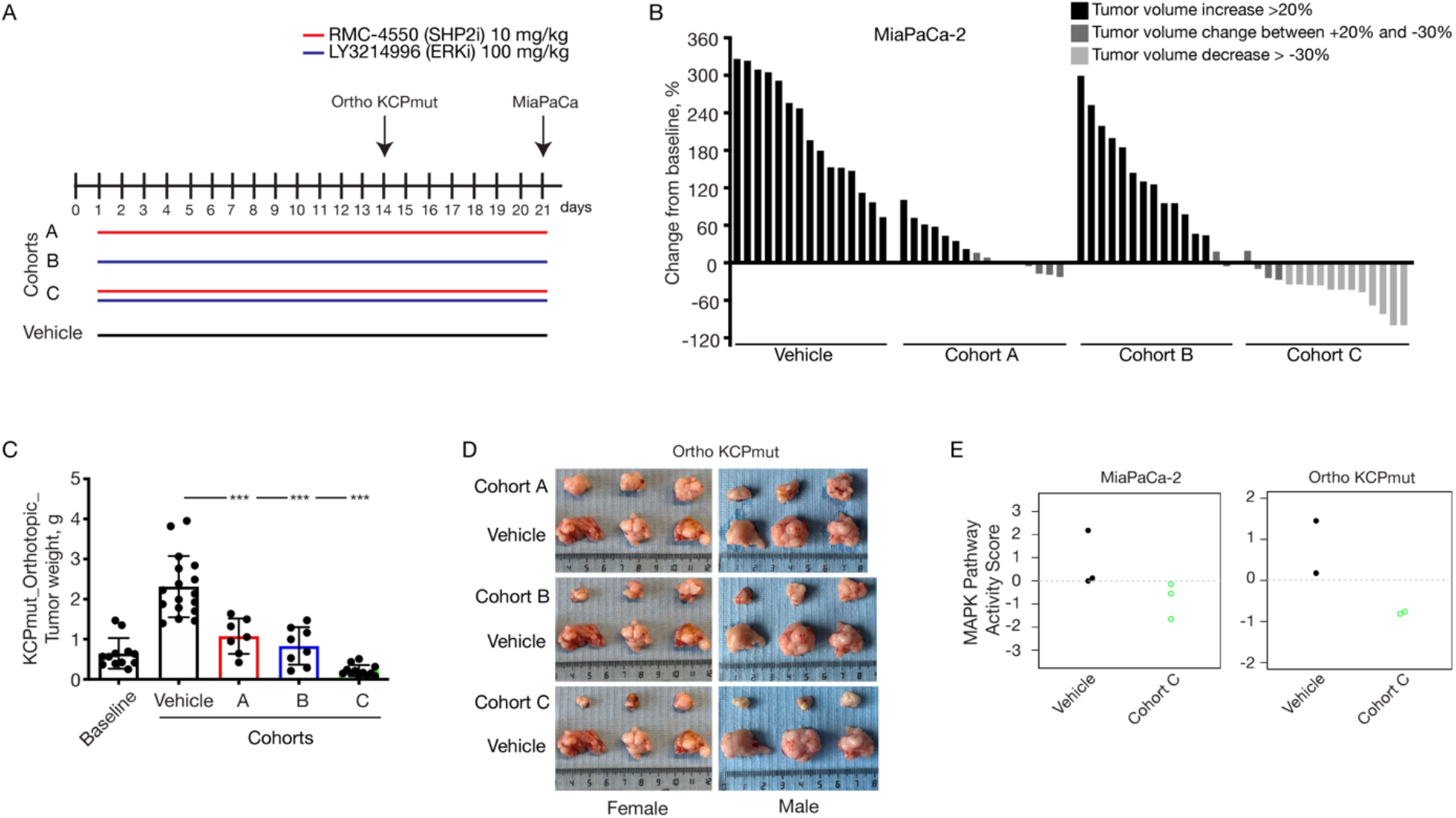
*In vivo* assessment of treatment response in a xenograft and in an orthotopic PDAC model. **A**: Schematic representation of the treatment schedule applied in a xenograft model of subcutaneously transplanted MiaPaCa-2 cell line and in a model of orthotopically transplanted KPC^mut^ tumors. Cohort A: Continuous treatment with RMC-4550 (SHP2i) alone daily; Cohort B: Continuous treatment with LY3214996 (ERKi) alone daily; Cohort C: Continuous treatment with the combination of SHP2i and ERKi daily. Control mice were continuously treated with vehicle. **B**: Evaluation of ERKi and SHP2i monotherapy treatments and combined administration of SHP2i and ERKi. For all the xenograft experiments, 5 × 10^6^ MiaPaCa-2 cells were subcutaneously injected into the right flank of NOD scid gamma (NSG) mice. When tumors reached 200 - 250 mm^3^, mice were randomly assigned into cohorts and treated by oral gavage with inhibitors or vehicle according to treatment schedule for 21 days, after which tumors were resected. The y axis shows tumor volume change in % from baseline. Each bar represents the difference in pancreatic volume in an individual animal. According to the RECIST criteria, black indicating progressive disease, dark grey indicating stable disease, and light grey indicating partial response. Significance was determined by one-way ANOVA with Bonferroni’s multiple comparison test. **C, D**: *In vivo* assessment of treatment response of orthotopically implanted tumors. ~40 mm^3^ tumor pieces (KCP^mut^) were orthotopically implanted into the pancreata of 8-week-old male and female C57BL/6 mice. After 2 weeks, mice were either sacrificed as baseline (n = 12) or randomly assigned into cohorts and treated with inhibitors or vehicle according to the treatment schedule (**A**). (**C**) Tumor weight (mean ± SD) was determined after 14 days of therapy as indicated: Baseline (n = 12), Vehicle (n = 17), Cohort A (n = 7), Cohort B (n = 8), Cohort C (n = 12). **** P < 0.0001, as determined by oneway ANOVA with Bonferroni’s multiple comparison test. (**D**) Representative macroscopic photographs of tumors in (**C**). **E**: MAPK pathway activity scores in MiaPaCa-2 xenograft and in KPC^mut^ orthotopic mouse models.

### Optimal treatment regimen

Having shown a potent anti-tumor benefit of the ERKi + SHP2i combination treatment, compared to the monotherapies in MiaPaCa-2 xenografts and orthotopic KCP^mut^ tumors, we decided to further expand our validation in additional PDAC models of murine and human origin, and at the same time compare intermittent treatment schedules. With regards to future clinical application and the possibility of adverse effects in patients, we wanted to determine the optimal treatment regimen, defined as maximum anti-tumor effect with minimum toxicity. For the ERKi + SHP2i combination these schedules included: continuous administration (daily) of both drugs in combination (Cohort C), as well as three different non-continuous (intermittent) schedules (Cohort D, E, and F). Control cohorts included daily vehicle treatment, as well as intermittent ERKi or SHP2i monotherapy (Cohort G, H, and I) (**Figure 4 A**).

**Figure 4.**
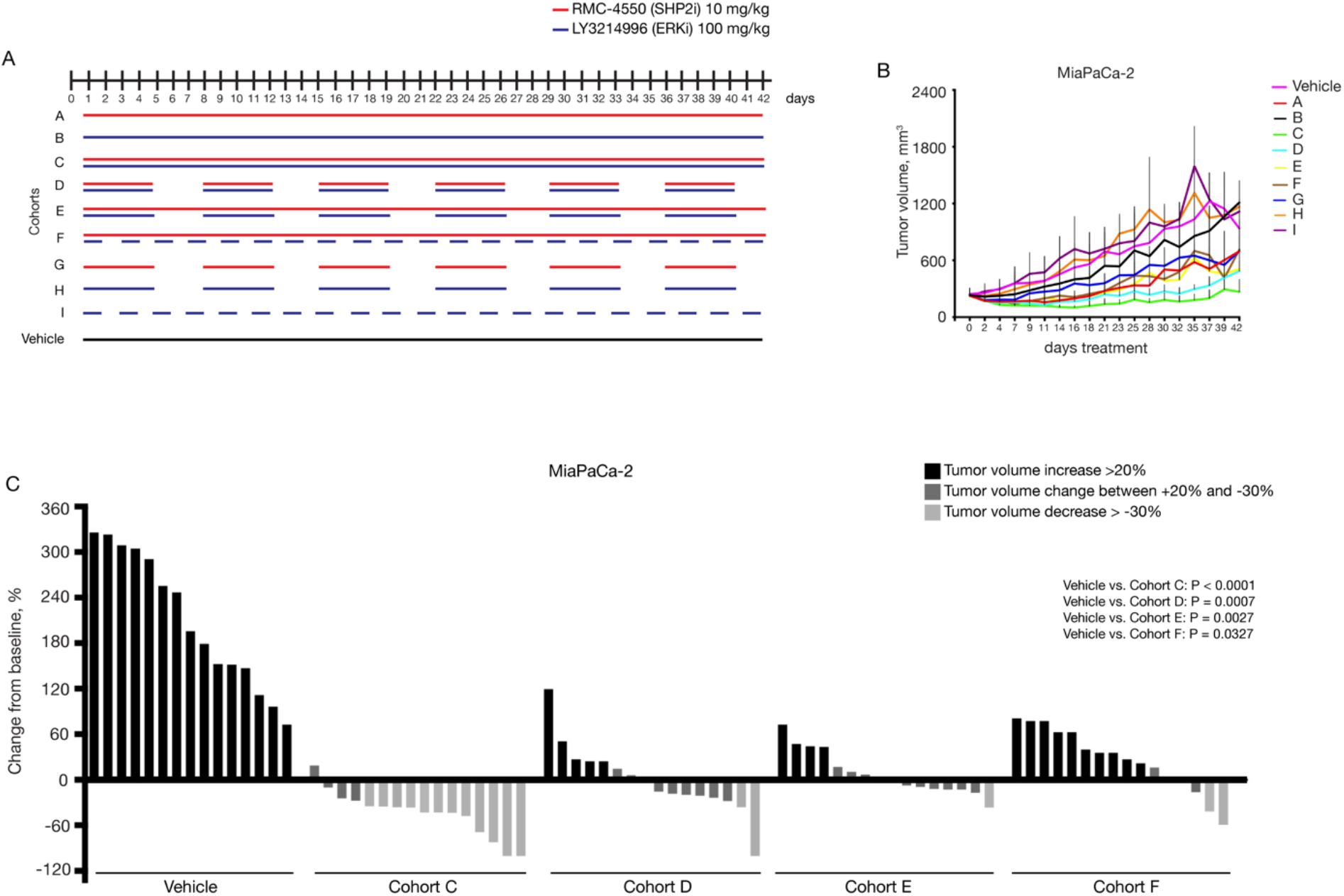
*In vivo* assessment of optimal treatment regimen in a xenograft model. **A**: Schematic representation of the treatment schedule applied in MiaPaCa-2 xenograft model. Cohort A: Continuous treatment with SHP2i alone daily; Cohort B: Continuous treatment with ERKi alone daily; Cohort C: Continuous treatment with the combination of SHP2i and ERKi daily; Cohort D: Intermittent treatment with the combination of SHP2i and ERKi 5 days on / 2 days off; Cohort E: Semi-continuous treatment schedule with daily dosing of SHP2i and intermittent dosing with ERKi 5 days on / 2 days off. Cohort F: Continuous treatment with SHP2i and on alternate days with ERKi. Cohort G: Intermittent dosing with SHP2i alone 5 days on / 2 days off. Cohort H: Intermittent dosing with ERKi alone 5 days on / 2 days off. Cohort I: Treatment with ERKi alone on alternate days. Control mice were continuously treated with vehicle. For all the xenograft experiments, 5 × 10^6^ cells were subcutaneously injected into the right flank of NOD scid gamma (NSG) mice. When tumors reached 200 - 250 mm^3^, mice were randomly assigned into cohorts and treated by oral gavage with inhibitors or vehicle according to treatment schedule. **B**: Treatment response was assessed through tumor volume change using caliper measurements 3 times/week in MiaPaCa-2 (KRAS^G12C^) xenograft model. Results represent mean ± SD. **C**: Tumor volume change at time point day 21. The y axis shows tumor volume change in percentage from baseline. Each bar represents the difference in tumor volume in an individual animal. According to the RECIST criteria, black indicating progressive disease, dark grey indicating stable disease, and light grey indicating partial response. Vehicle and Cohort C data from **Figure 3 B** are reported again for comparison. Significance was determined by one-way ANOVA with Bonferroni’s multiple comparison test.

We tested these regimens in three different tumor models: the subcutaneous xenograft model with transplanted human PDAC cell lines (**Figure 4 and Supplementary Figure S2**), the endogenous *Kras; Trp53^-/-^* (KCP) model of spontaneous PDAC formation (**Figure 5 and Supplementary Figure S3**), and the subcutaneous model of transplanted patient derived PDAC tissue xenografts (PDX) (**Figure 6 and Supplementary Figure S4**). None of the tested schedules were associated with dose or schedule-limiting toxicities (**Supplementary Figure S5**) in either of the three tumor models.

**Figure 5.**
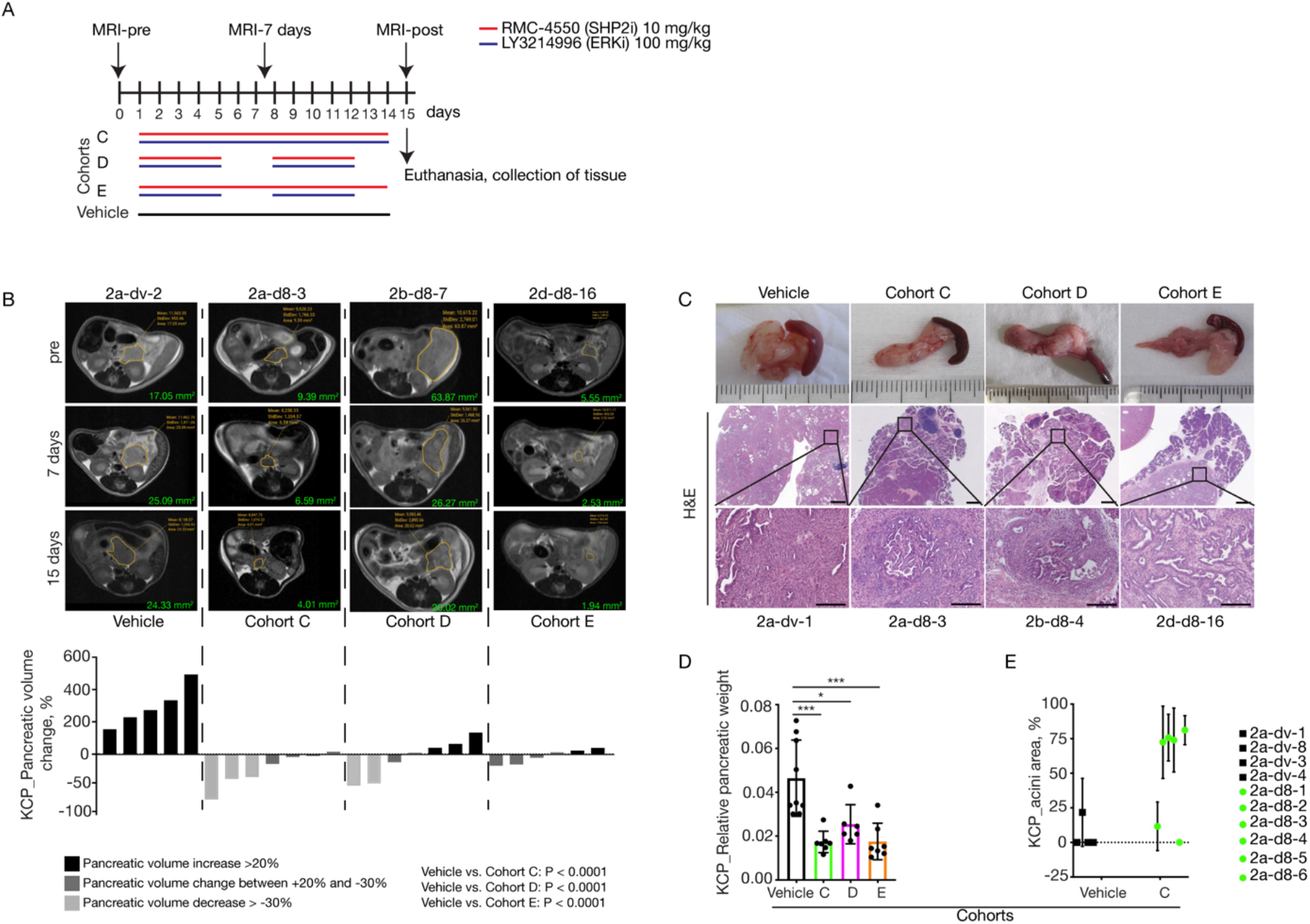
*In vivo* assessment of optimal treatment regimen in an endogenous murine PDAC model. **A**: Schematic representation of the treatment schedule applied in the endogenous (KPC) murine model of spontaneous tumor formation as well as the Magnetic Resonance Imaging (MRI) time points applied. Cohort C: Continuous treatment with the combination of SHP2i and ERKi daily; Cohort D: Intermittent treatment with the combination of SHP2i and ERKi 5 days on / 2 days off; Cohort E: Semi-continuous treatment schedule with daily dosing of SHP2i and intermittent dosing with ERKi 5 days on / 2 days off. Control mice were treated with vehicle for 14 consecutive days. All treated mice were sacrificed on day 15, and tumors were resected for histological analysis. **B**: Representative MRI scan slices depicting PDAC tumor sections of KCP mice treated with vehicle (n = 5), Cohort C (n = 7), Cohort D (n = 7) or Cohort E (n = 6) at the indicated time points (days) following the start of therapy (pre), with similar results among the groups. Volumetric measurements indicate a decrease in pancreatic volume in mice treated with the combination of SHP2i and ERKi for two weeks compared to vehicle-treated mice. The y axis shows pancreatic volume change in % quantified by measurements of MRI scans. Each bar represents the difference in pancreatic volume in an individual animal from day 0 to day 15. According to the RECIST criteria, black indicating progressive disease, dark grey indicating stable disease, and light grey indicating partial response. Significance was determined by one-way ANOVA with Bonferroni’s multiple comparison test. Volume-tracking curves for individual mice over the whole course of therapy are available in **Supplementary Figure 3 B-E**. **C**: Macroscopic images of pancreas and spleen (top row). Representative H&E-stained sections of pancreata from mice, treated as indicated. Scale bars represent 1000 μm (middle) and 200 μm (bottom). Mice numbers are indicated below. **D**: Relative pancreatic weight was significantly lower in all groups treated with the combination of SHP2i and ERKi: Cohort C (n = 7); Cohort D (n = 6); Cohort E (n = 7) compared to vehicle-treated control mice (n = 9). Results represent mean ± SD. * P < 0.05, *** P < 0.001, significance was determined by one-way ANOVA with Bonferroni’s multiple comparison test. **E**: Quantification of relative intact acinar area as ratio of whole pancreatic area. Analysis performed on n = 4 individual mice in vehicle group and on n = 6 individual mice in Cohort C.

**Figure 6.**
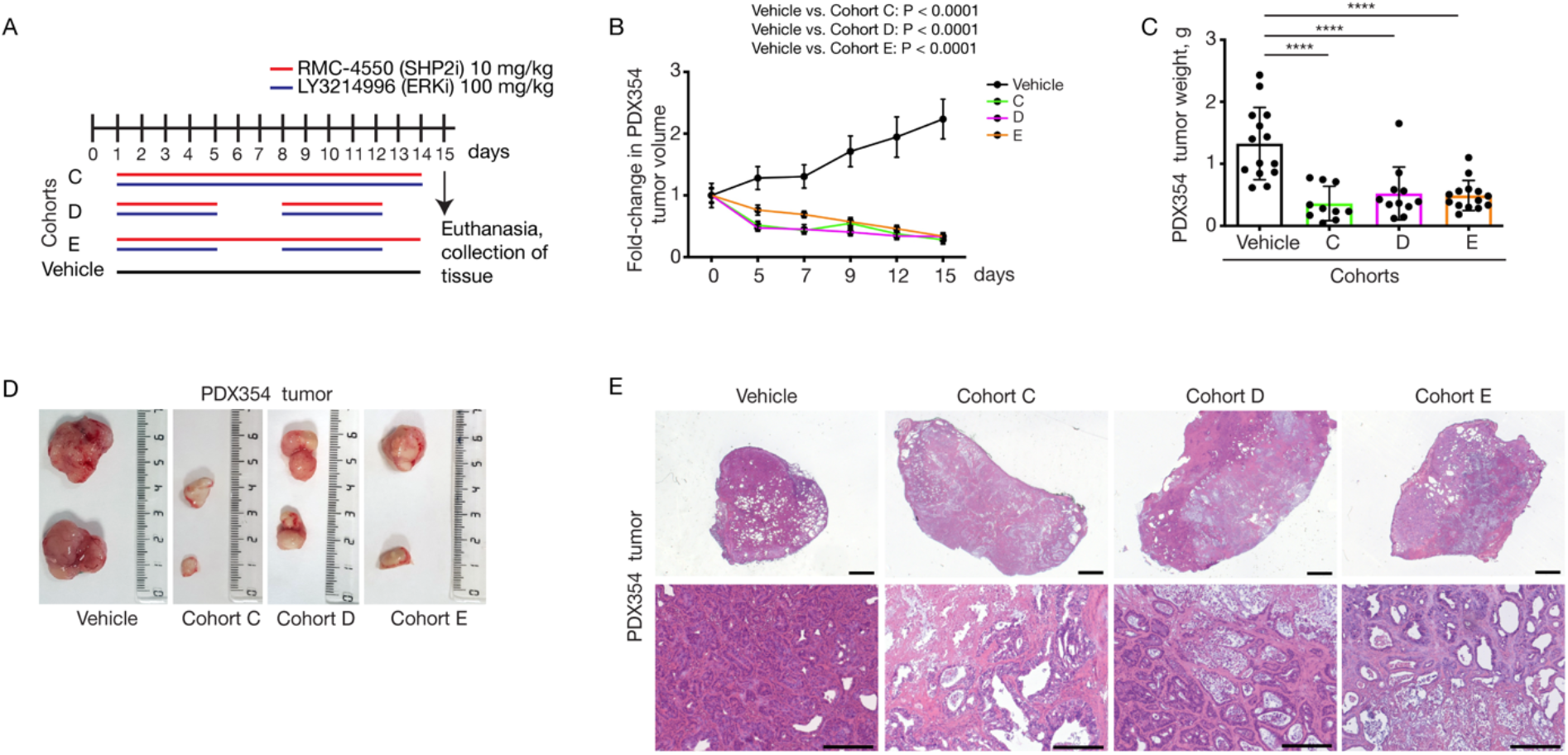
Evaluation of treatment response by the combined administration of RMC-4550 (SHP2i) and LY3214996 (ERKi) in Patient-Derived Xenograft (PDX) models. **A**: Treatment schedule. Mice were treated with the combination of RMC-4550 (SHP2i) and LY3214996 (ERKi) once per day via oral gavage for 14 consecutive days (Cohort C) or 5 days on/2 days off (Cohort D) or SHP2i continuously and ERKi 5 days on / 2 days off (Cohort E). For PDX354 model, tumor pieces of 50 mm^3^ were subcutaneously implanted into both flanks of NSG mice. When tumors reached 200 - 250 mm^3^ (approximately 6 - 8 weeks after subcutaneous transplantation), mice were randomly assigned into cohorts and treated by oral gavage with inhibitors or vehicle according to treatment schedule for the indicated time, after which tumors were resected. **B, C**: Treatment response was assessed through tumor volume changes using daily caliper measurements (**B**) and tumor weight at endpoint (**C**) in PDX354 model. Results represent mean ± SD. **** P < 0.0001, as determined by one-way ANOVA with Bonferroni’s multiple comparison test. **D**: Representative macroscopic images of resected tumors. **E**: Representative H&E-stained sections of vehicle- and combination therapy-treated PDX354 tumors. Scale bars represent 1000 μm (top) and 200 μm (bottom).

The potent tumor inhibitory effect of the continuous combination treatment (Cohort C) shown in MiaPaCa-2 xenografts (**Figure 4 C**) was also observed in Panc 10.05, ASPC1, and YAPC xenografts (**Supplementary Figure S2**). While we observed varying degrees of sensitivity amongst the 4 PDAC cell lines, the common findings were that all combination schedules showed stronger anti-tumor efficacy compared to all monotherapy or vehicle controls, and that the continuous schedule had the strongest inhibitory effect of all the combination regimens (**Figure 4 C and Supplementary Figure S2**). However, complete tumor elimination was not achieved, even with the continuous treatment schedule.

In agreement with these findings, our endogenous KCP model showed that the combination treatment was able to significantly inhibit tumor growth (**Figure 5 A-D**) and even induce pancreatic volume reduction. Of note, the continuous treatment was able to induce more than 30% pancreatic volume reduction (**Figure 5 B**) and prevent tumor outgrowth from the microscopic to macroscopic scale if the mouse was treated early enough. This was indicated by morphological analysis, the relative pancreatic weight, and the considerable number of intact acini compared to the control (**Figure 5 C-E**).

Additionally, three different PDX models, each representing a different patient harboring KRAS^G12D^ mutations, corroborated the observation that all tested combination therapy regimens were able to significantly reduce tumor growth and even induce tumor volume reduction (**Figure 6 and Supplementary Figure S4**). While there was no statistically significant difference between the three treatment arms, the data seem to indicate a slight benefit of the continuous treatment compared to the intermittent schedules (**Figure 6 C-D and Supplementary Figure S4 C and G).**Interestingly, all tumors from mice receiving the combination treatments contained lytic necrotic cores, suggesting tumor cell elimination in addition to the cytostatic affects observed (**Figure 6 E and Supplementary Figure S4 D and H).**

### Assessment of treatment response

Having shown efficacy data in multiple different *KRAS*-mutant PDAC mouse models, our aim is to bring this novel therapy to the clinic. While we have tried to diversify our models and thus account for inter-patient heterogeneity, it is still essential to distinguish responders from nonresponders in a clinical setting, preferably using minimally invasive methods. Interestingly, Ying et al. (10) described that oncogenic KRAS^G12D^ is required for PDAC tumor maintenance and reprograms PDAC metabolism by stimulating glucose uptake and glycolysis, while Bryant et al. (45) found that both KRAS suppression and ERK inhibition decreased glucose uptake in PDAC. Based on those reports, we explored the possibility of using ^18^F-FDG (fluorodeoxyglucose) uptake, as measured by PET scan, as early response marker and a surrogate readout of MAPK activity.

To this aim, we treated mice bearing subcutaneous KCP tumors with either vehicle or the combination of RMC-4550 + LY3214996 daily for 7 days. PET scans were performed at days 0 (pre-treatment), 3 and 7. Our results show that ^18^F-FDG uptake is readily detected by PET-CT scan of subcutaneously implanted KCP tumors, and that a significant decrease in PET signal is already observable at a time when reduction in tumor volume is not yet significant (**Figure 7**). This finding raises the possibility of using FDG uptake in the clinic to monitor early response to this novel combinatorial therapy for PDAC patients.

**Figure 7.**
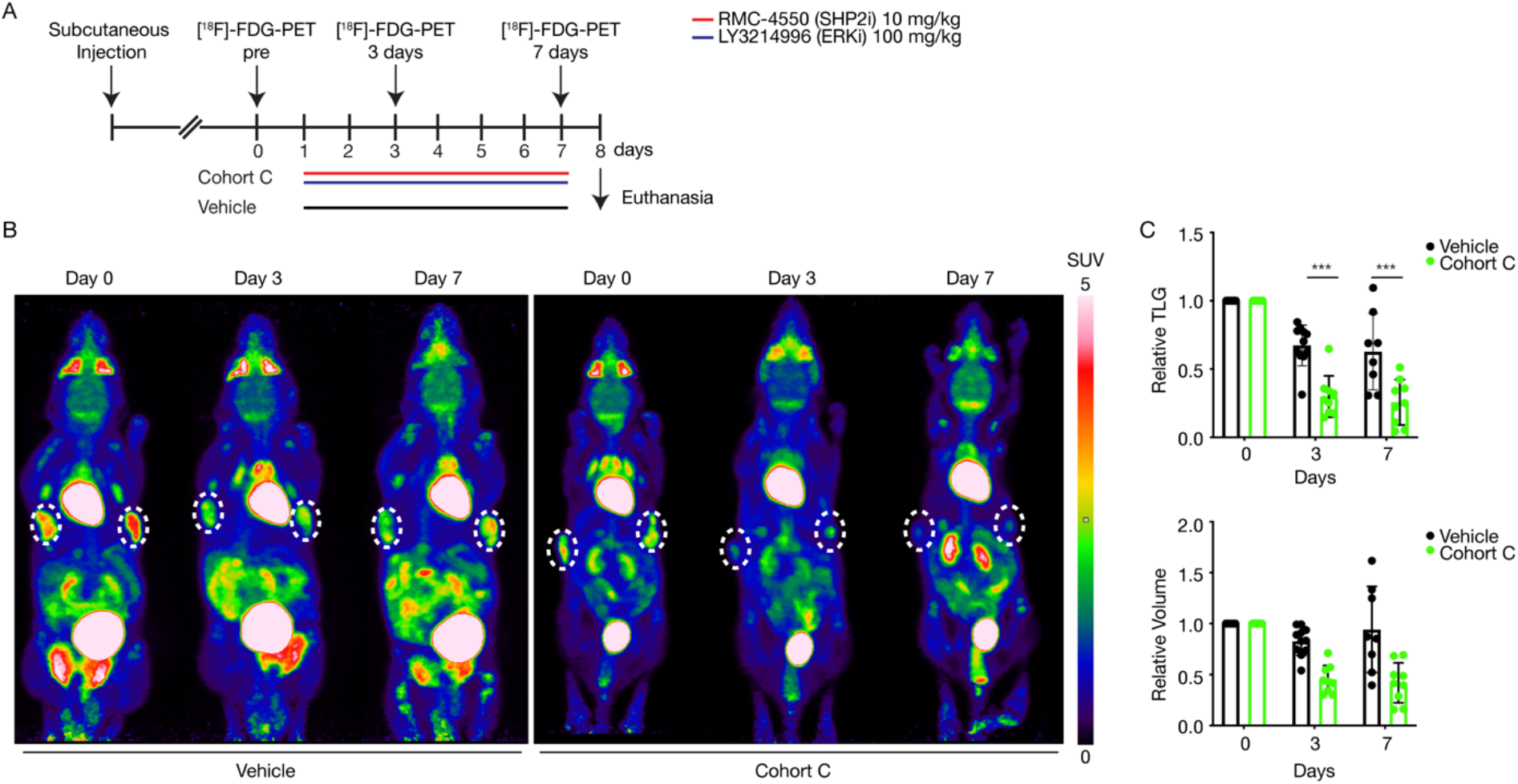
Early non-invasive assessment of the treatment response in a subcutaneous tumor mouse model. **A**: Schematic representation of the treatment schedule applied in the subcutaneous tumor mouse model as well as the ^18^F-FDG-PET imaging time points applied. 2.5 × 10^6^ - 3 × 10^6^cells were injected subcutaneously into the left and right flank of 10- to 15-week-old non-tumor-bearing littermates. 2 - 3 weeks after subcutaneous injection, mice were randomly assigned into cohorts and treated by oral gavage with inhibitors or vehicle for 7 consecutive days. ^18^F-FDG-PET scans were obtained at baseline before commencement of therapy (day 0) and at days 3 and 7 during treatment. **B:** Representative ^18^F-FDG-PET images of tumor bearing mice treated with vehicle or undergoing treatment with the combination of RMC-4550 (SHP2i) and LY3214996 (ERKi) once per day for 7 consecutive days (Cohort C). Subcutaneous tumor areas are shown in dashed circles, SUV of FDG uptake is indicated by color. White color indicates highest uptake, red color - high uptake, yellow - green (intermediate), blue represents low uptake. ^18^F-FDG-PET = 18-fluordesoxyglucose Positron Emission Tomography; PET = Positron Emission Tomography; SUV = Standardized Uptake Values; FDG = Fluorodeoxyglucose. **C**: Relative Total Lesion Glycolysis (TLG) on days 0, 3, and 7 in Vehicle vs. Cohort C (n = 8 - 9 animals/group). **C**: Relative tumor volume on days 0, 3, and 7 in Vehicle vs. Cohort C. Results represent mean ± SD. *** P < 0.001, significance was determined by unpaired, two-tailed t test.

## Discussion

Pancreatic cancer has one of the highest mortality rates amongst all tumor types, and innovative treatment options against this devastating cancer are urgently needed (2). Since targeted therapies are lacking and PDAC seems to be refractory to immunotherapy (46), classic chemotherapy is still the treatment of choice for the management of PDAC at all stages of the disease (47).

We recently identified a novel strategy for targeting *KRAS*-mutant tumors, irrespective of the specific mutation, which consists of the concomitant blockade of the RAS downstream effector MEK and the upstream activator SHP2 (25, 27). Similarly, the combination of SOS1 and MEK inhibition has been proven effective in *KRAS*-mutant preclinical tumor models (48). In the present work, we validate this “up plus down” double blockade strategy in multiple *in vivo* models of PDAC by combining inhibition of the MEK downstream effector ERK (by using LY3214996), with upstream inhibition of SHP2 (with RMC-4550).

In agreement with our earlier findings, we show that ERK inhibitor monotherapy is insufficient to induce a durable suppression of the MAPK pathway, as demonstrated by a rebound in phosphorylated RSK1 levels and failure to induce apoptosis. We have previously demonstrated that the rebound in MAPK signaling following RAS downstream inhibition can be attributed to the feedback overexpression and activation of multiple receptor tyrosine kinases. Therefore, the most effective way to short-circuit this resistance-inducing loop is to disrupt the signal transmission from activated RTKs to RAS. The protein phosphatase SHP2 has been found to be recruited by virtually all phosphorylated growth factor receptors, as well as other cell surface receptors like cytokines or hormone receptors, where it mediates the signal transmission to downstream protein effectors, making it the ideal target to prevent ERK inhibitor resistance (49). Indeed, we show that co-treatment with LY3214996 and RMC-4550 promptly induces apoptosis in cell cultures and tumor regression in several mouse models of PDAC, especially when administered continuously.

The previous failure of MEK inhibitors like selumetinib against *KRAS*-mutant tumors during clinical trials (50) has been attributed not only to the above-mentioned resistance mechanisms (22), but also to the highly toxic profile of such drugs. On the other hand, ERK inhibitors are novel compounds that only recently have entered the earliest phases of clinical testing, holding promise for improved tolerability and efficacy in the treatment of tumors with a MAPK pathway dependency. In the present study, we extensively evaluated the toxicity of the LY3214996 + RMC-4550 combination in multiple murine backgrounds, and identified for each compound a dose that, in combination, is well tolerated as well as efficacious against PDAC tumor growth.

Apart from its role in the MAPK pathway, the SHP2 phosphatase is also recruited by the immune checkpoint receptor PD1, which is expressed on T and pro-B lymphocytes (51). PD1 activity is known to suppress T cell activation and therefore mediate cancer immune evasion (52, 53). It has been reported that inhibition of SHP2 can stimulate an anti-tumor immune response by both promoting T-cell function and depleting pro-tumorigenic M2 macrophages (54, 55). Therefore, the use of SHP2 inhibitors could have a double beneficial effect: in the tumor cells it would act synergistically with ERK inhibitors to suppress MAPK-induced proliferation, and in the tumor-microenvironment it could promote the anti-tumor immune response. Further investigation, for example using a pancreatic cancer model that can be transplanted both in immuno-competent and immuno-compromised hosts, will be needed in order to elucidate the non-tumor-intrinsic benefits of using SHP2 inhibitors.

As a translational approach to monitor the early response of patients, we searched for dynamic biomarkers that could be used in a clinical setting. So far, computed tomography is the method of choice to determine therapy response; however, this is usually done retrospectively. Thus, the idea of non-invasive but quick methods to evaluate an early drug response is gaining increasing attention in modern oncology, as it allows monitoring of therapy failure sparing the patient unnecessary toxicity. In 2006, Su et al., (56) showed that inhibition of the MAPK pathway by treatment with anti-EGFR therapy in colorectal cancer models induced a rapid downregulation of glucose receptors, which could be reflected by decreased FDG glucose uptake in PET scans. Recently, Bryant et al. (45) reported that siRNA-mediated silencing of KRAS or pharmacological ERK inhibition decreased glucose uptake and resulted in a clear reduction of key glycolytic intermediates in KRAS-driven PDAC. Furthermore, a pilot study by Wang et al. (57) showed that ^18^F-FDG-PET and diffusion weighted MRI (DW-MRI) can be used as early treatment response assessment in patients with advanced PDAC. Based on these data, we suggest ^18^F-FDG imaging via PET scans as a tool to sensitively monitor tumor shrinkage shortly after therapy initiation to evaluate the early drug response. Indeed, we confirmed that co-inhibition of ERK and SHP2 in subcutaneous *KRAS*-mutant PDAC tumors induces a rapid and significant decrease in FDG uptake, which precedes the decrease in tumor volume.

In conclusion, LY3214996 + RMC-4550 combination has shown a positive tolerability profile as well as the capacity to synergistically induce a high percentage of partial response in several preclinical models of PDAC. In addition, the preliminary data of the PET scans to monitor early drug response, warrants the ERK + SHP2 inhibitor combination to be explored at a clinical level. To this end, we started the Phase 1a/1b SHERPA trial (NCT04916236) with the objective of testing the tolerability and early evidence of efficacy of the combination of RMC-4630 (the clinical equivalent of RMC-4550) and LY3214996, which may represent a promising targeted therapeutic option for the majority of *KRAS*-mutant pancreatic cancer patients.

## Methods

### Mouse strains

*Kras^G12D^* (*Kras^tm4Tyj^*) (58); *p48-Cre* (*Ptf1a^tm1(Cre)Hnak^*) (59); *p53^flox/flox^(Trp53^tm1Brn^*) (60) (KCP) have been described previously and were bred in a mixed genetic background in our animal facility. Non-tumor-bearing littermates without mutational *Kras^G12D^* (*Kras^tm4Tyj^*) and *p48-Cre* (*Ptf1a^tm1(Cre)Hnak^*) were used in the dose finding study and for subcutaneous tumor transplantation. *Kras^G12D^* (*Kras^tm4Tyj^*); *Pdx1-Cre* (*Tg(Pdx1-cre)6Tuv*); *Trp53^mut/+^(Trp53^tm1Tyj^*) (61) (KCP^mut^) mouse strain with C57BL/6J background served as tumor donor for orthotopic transplantation experiments.

At the age of weaning and after death, genotypes were determined by PCR and gel electrophoresis. B6 (C57BL/6J) mice and NSG (*NOD.Cg-Prkdc^scid^ II2rg^tm1Wjl^/SzJ*) mice were obtained from Jackson Laboratory. NU-Foxn1nu nude mice (*Hsd:Athymic Nude-Foxn1^nu^/Foxn1^+^*) were obtained from Envigo.

All mice were kept in an animal room (room temperature range between 20 and 22°C) with light-dark cycle of 12:12 hours in groups of 2–4 animals in type III cages (Tecniplast) or in groups of 5 animals in IVC cages from (Innovive) with bedding and nesting material. All animals were provided with the standard maintenance food for mice (No. 1324 – 10 mm pellets, Altromin, or SDS diets Technilab BMI) and water ad libitum and housed under specific pathogen-free conditions in accordance with the European Directive 2012/63/EU.

### Cell culture and cell lines

Primary murine tumor cell lines were obtained from chopped pieces of explanted tumors without enzymatic digestion. All murine cell lines were routinely cultured in DMEM supplemented with 10% FBS and penicillin–streptomycin (100 U/ml, 100 μg/ml) (all Life Technologies). Human PDAC cell lines YAPC (*KRAS^p.G12V^*; *p53^p.H179R^*; *SMAD4^p.R515fs^*^22^*), ASPC1 (*KRAS^p.G12D^*; *p53^p.C135fs*35^*; *SMAD4^p.R100T^*; *CDKN2A^p.L78fs*41^*), Panc 10.05 (*KRAS^p.G12D^*; *p53^p.1255N^*), Panc 1 (*KRAS^p.G12D^*; *p53^p.R273H^*) and MiaPaCa-2 (*KRAS^p.G12C^*; *p53^p.R248W^*; Homozygous for *CDKN2A* deletion) were purchased from the American Type Culture Collection (ATCC). Mutational status of the cell lines was compiled from the ATCC, Catalogue of Somatic Mutations in Cancer (COSMIC; Wellcome Trust Sanger Institute) and Cancer Cell Line Encyclopedia (CCLE, Broad Institute) databases. Human cell lines were cultured in RPMI1640 (Life Technologies), supplemented with 10% FBS, penicillin/streptomycin (100 U/ml, 100 μg/ml, Life Technologies), and 2mM L-glutamine (Thermo Fisher Scientific). All cells were kept at 37 °C in a humidified incubator with 5% CO_2_.

### Drugs and inhibitors

SHP2 inhibitor RMC-4550 (SHP2i) was kindly provided by Revolution Medicines, Redwood City, California U.S.A. SHP2i was diluted in 50 mM Sodium Citrate Buffer pH = 4 with 1% Hydroxyethylcellulose (Sigma-Aldrich), 0.25% Tween (Sigma-Aldrich) and 0.05% Antifoam A concentrate (Sigma-Aldrich). Erk1/2 inhibitor LY3214996 (ERKi) was kindly provided by Eli Lilly and Company, Indianapolis IN 46285 U.S.A. ERKi powder was dissolved in dH_2_O (Braun) with 1% Hydroxyethylcellulose (Sigma-Aldrich), 0.25% Tween (Sigma-Aldrich) and 0.05% Antifoam A concentrate (Sigma-Aldrich). Inhibitor combinations were used according to company’s recommendation.

### *In vitro* drug synergy and quantitative analysis

Indicated cells were cultured and seeded into 96-well plates at a density of 300–2,000 cells per well, depending on growth rate. Twenty-four hours later, drugs were added at the indicated concentrations using the HP D300 Digital Dispenser (HP). After 72 hours, medium and drugs were refreshed. The total duration of the experiment was 6 days (two treatments) for KCP_K2101, KCP_P0012, MiaPaCa-2 and Panc10.05, and 10 days (three treatments) for Panc1, ASPC1 and YAPC. Cells were fixed with 4% PFA diluted in PBS (37% Formaldehyde solution, Merck) and stained with 2% crystal violet solution (HT90132-1 L, Sigma-Aldrich). Drug synergy was calculated using CompuSyn software (version 1.0), which is based on the median-effect principle and the combination index–isobologram theorem (62). CompuSyn software generates combination index values, where combination index 0-0.75 indicates synergy, 0.75-1.25 indicates an additive effect and CI >1.25 indicates antagonism (63). Following the instructions of the software, drug combinations at non-constant ratios were used to calculate the combination index in our study.

### Incucyte cell-proliferation assay and apoptosis assay

Indicated cell lines were seeded into 96-well plates at a density of 200–2,000 cells per well, depending on growth rate and the design of the experiment. Approximately 24 hours later, drugs were added at the indicated concentrations using the HP D300 Digital Dispenser (HP). Cells were imaged every 4 hours using the Incucyte ZOOM (Essen Bioscience). Phase-contrast images were analyzed to detect cell proliferation on the basis of cell confluence. For cell apoptosis, caspase-3 /caspase-7 green apoptosis-assay reagent (Essen Bioscience) was added to the culture medium (1:1000), and cell apoptosis was analyzed on the basis of green-fluorescent staining of apoptotic cells.

### *In vivo* drug combination dose finding escalation

Dose finding was established according to modified “3 + 3” scheme. Non-tumor-bearing mice were put on continuous oral administration of both drugs over 14 days (NSG mice, *Kras^G12D^*; *p53^flox/flox^*, *p53^flox/flox^*, *p53^flox/wt^* or wildtype mice from *Kras^G12D^*; *p48-Cre*; *p53^flox/flox^* litter). Toxicity was evaluated daily by measuring mice body weight (endpoint at body weight loss > 20%), general clinical signs (abnormal behavior, signs of physical discomfort). According to modified “3 + 3” design, mouse cohorts consisting of 3 animals were given an initial combination dose (d5), followed by increased dose 7 as no side effects were observed in all 3 mice. Up to six mice were assigned to one dose. If the combination dose showed side effects in 1/6 mice, the dose was designated as an admissible dose, opening next dose level for testing. If dose-limiting toxicity was observed in 2/6 mice, the combination was accepted as a maximum tolerated dose, closing higher doses for testing. If more than two of the six mice experienced dose-limiting toxicity, the dose was down staged. The following dose combinations were administered: dose 5 (10 mg/kg RMC-4550 + 75 mg/kg LY3214996), dose 7 (30 mg/kg RMC-4550 + 75 mg/kg LY3214996), dose 8 (10 mg/kg RMC-4550 + 100 mg/kg LY3214996) and dose 9 (30 mg/kg RMC-4550 + 100 mg/kg LY3214996). One cohort was administered vehicle (50 mM Sodium Citrate Buffer pH =4 with 1% Hydroxyethylcellulose, 0.25% Tween and 0.05% Antifoam A concentrate) to monitor gavage-mediated side effects.

### *In vivo* therapy treatment schedules

For *in vivo* application in KCP mice, human cell line xenografts, PDX xenografts, orthotopically transplanted KCP^mut^ mice, and subcutaneously transplanted KCP mice, dose 8 (10 mg/kg RMC-4550 + 100 mg/kg LY3214996) was administered by oral gavage. Depending on the mouse model, the drug administration followed different schedules: continuous administration (daily) of both drugs (cohort C), administration of both drugs 5 days on/2 days off (cohort D), or continuous administration of RMC-4550 plus LY3214996 every other day (cohort E), or continuous administration of RMC-4550 plus LY3214996 5 days on/2 days off (cohort F). As controls, mice were treated daily with vehicle or with monotherapy. Monotherapy was scheduled either RMC-4550 continuous (cohort A) or 5 days on/2 days off (cohort G). Monotherapy with LY3214996 was accomplished continuously (cohort B), 5 days on/2 days off (cohort H) or every other day (cohort I).

### Patient-derived tissue xenografts

Previously established PDAC (all KRAS^G12D^) patient-derived xenografts (PDAC PDX) were obtained from Dr. Manuel Hidalgo under a Material Transfer Agreement with the Spanish National Cancer Research Center (CNIO), Madrid, Spain (Reference no. I409181220BSMH). The indicated PDXs were expanded in female 6- to 8-week-old NU-Foxn1nu nude mice (Envigo). For subcutaneous transplantation of patient-derived tumor tissue, a single ~ 50 mm^3^ section of tumor embedded in Matrigel (Corning) was implanted into subcutaneous pockets in the posterior flanks of 6-week-old NSG male mice. Volumes were evaluated every 2 days by caliper measurements and the approximate volume (V) of the mass was estimated using the formula V = D × d^2^ / 2, with D being the major tumor axis and d being the minor tumor axis. According to tumor volume (average volume 50 – 150 mm^3^) treatment was initiated 6 - 8 weeks after surgery. Mice were randomly assigned to trial arms (cohorts C, D, E and Vehicle). Experiments were terminated once vehicle control tumors reached a critical size at the ethical end point (V = 2000 mm^3^). End-of-treatment tumor material was formalin-paraffin embedded.

### Human pancreatic cancer cell line xenografts

MiaPaCa-2, Panc 10.05, ASPC1, and YAPC cells were resuspended (5 × 10^6^ cells per mouse) in a 1:1 mixture of RPMI and Matrigel (Corning) and injected subcutaneously into the right flanks of 8-week-old NSG mice. Tumor volume was monitored three times a week as described for the patient-derived tissue xenografts. Mice were randomized when the tumor reached a volume of approximately 200 mm^3^ and treated for a maximum period of 30 days (YAPC) or 42 days (MiaPaCa-2, Panc 10.05, ASPC1). Mice were sacrificed after 1, 3 or 6 weeks of treatment (8 mice per time point and per cohort) or at humane end point. In this experimental set up, RMC-4550 (10 mg/kg) and LY3214996 (100 mg/ kg) were dissolved in 2% HPMC E-50, 0.5% Tween-80 in 50 mM Sodium Citrate Buffer, pH 4.0, and administered according to the different schedules (cohorts A, B, C, D, E, F, G, H, I). Control groups were treated daily with the vehicle alone. End-of-treatment tumor material was partly snap frozen in liquid nitrogen and stored at −80 °C.

### Orthotopic PDAC mouse models

30 mm^3^ KCP^mut^ tumor pieces were obtained from endogenous mouse models and subcutaneously transplanted into the flanks of female B6 host mice for expansion. Tumor growth was monitored as indicated for the patient-derived tissue xenografts by caliper analysis. After 4 weeks, subcutaneous KCP^mut^ tumors from donor mice were harvested and chopped into ~ 40 mm^3^ pieces and orthotopically transplanted into pancreata of 8-week-old female and weight matched 18 – 20 g male B6 mice, as previously described (64). Tumor growth was monitored by palpation. After 2 weeks, a representative number of mice were sacrificed to determine pre-treatment baseline pancreas/tumor weights, and the remaining mice were randomly grouped into cohort A, B, C and vehicle and treated with inhibitors as described in the following. End-of-treatment tumor material was partly snap frozen in liquid nitrogen and stored at − 80 °C.

### Subcutaneous cancer cell line mouse models

Pancreatic cancer cells from KCP endogenous donor mouse model were obtained and cultured as described above. 2.5 × 10^6^ – 3 × 10^6^ cells were suspended in 100 μl of a 1:1 mixture of DMEM and Matrigel (Corning) and injected subcutaneously into the left and right flank of 10 - 15-week-old non-tumor-bearing female and male littermates from mixed background mouse strain *Kras^G12D^* (*Kras^tm4Tyj^*), *p53^flox/flox^* (*Trp53^tm1Brn^*). Tumor volume was monitored as indicated for the patient-derived tissue xenografts. Randomized therapy was initiated after tumors had reached a palpable volume of < 300 mm^3^. Therefore, female, and male mice were treated continuously with inhibitors (cohort C) or vehicle alone. On day 0, 3 and 7 of treatment, mice were scanned by animal PET (Mediso), imaging radioactive labelled glucose (^18^F-FDG) uptake. Mice were sacrificed after different time points, with a minimum treatment time of 3 days.

### PET imaging and ^18^F-FDG *in vivo*

Two-to-six hours fasted female and male mice bearing subcutaneous KCP tumors were randomly divided into two groups (vehicle versus cohort C). For PET imaging, mice received 12 – 14 MBq of the radiotracer ^18^F-Fluordeoxyglucose (18F-FDG) via injection through the lateral tail vein. PET images were acquired on a nanoScan PET system (Mediso, Budapest, Hungary) from 45 – 60 min post injection under isoflurane anesthesia (1 - 2 % in medical air by precision vaporizer (Baxter Healthcare, Deerfield, IL, USA). During the imaging procedure, mice were placed on a heated bed and their heart rate was constantly monitored. For 3D whole body image reconstruction with a 0.4 mm^3^ voxel size, the Tera-Tomo 3D image reconstruction algorithm (integrated into Nucline NanoScan Software, Mediso) was applied (4 iterations, 6 subsets), without AC and scatter corrections. Image counts per voxel per second were converted into standardized uptake values (SUV) using the activity concentrations computed the Nucline NanoScan software normalized to the animal’s body weight. Quantification of tumor uptake was carried out using the Nucline NanoScan software, by drawing spherical regions of interest (ROIs), creating a volume that represented the entire tumor lesion. We recorded the volume, SUVmean, SUVmax and total lesion glycolysis (TLG) of each tumor.

### Magnetic resonance imaging (MRI)

MR imaging for KCP male and female mice was started at an age of 24 - 37 days and repeated after 7 and 14 days of treatment (cohort C, D, E versus vehicle). Sedation was achieved via continuous inhalation of 2% isoflurane (Abbott) in 1.6% O_2_ using a veterinary anesthesia system (Vetland Medical). Body temperature was maintained and monitored, and eyes were protected by eye ointment. Image acquisition was achieved using a mouse 3T coil inside a preclinical 3T nano scan PET/MR (Mediso) and a T2 weighted fast spin echo sequence (resolution: 192 × 128 ~25 slices, echo time 55,52 ms; repetition time 3000 ms). Analysis, visualization, and calculation was done by Flywheel DICOM viewer. Solid tumor volumes were calculated by summating truncated pyramid volumes between tumor areas on vicinal slices. As drug treatment prevented tumor development in some of the endogenous KCP mice, pancreatic areas including tumor and non-neoplastic tissue were defined as regions of interest and summarized in the scanned slices to calculate pancreatic volume.

### Histology

Tissue specimen were fixed in 4% buffered paraformaldehyde for 48 hours at 4 °C, dehydrated and embedded in paraffin. H&E was performed as described previously on 2.5 μm cut sections (65). Image acquisition was performed on a Zeiss AxioImager.A1 microscope. Quantitative analyses of tumor and acini areas were performed with Axiovision (Zeiss).

### RNA sequencing and MAPK Pathway Activity Score

For preparation of the MiaPaCa-2 and the Panc 10.05 xenograft tumors, snap frozen material (3 mice from cohort C and vehicle, treated for 3 weeks), was cut at the cryostate (30 cryosections of 30 μm thickness per sample) and RNA was extracted using the RNeasy Mini Kit from Qiagen and analyzed using an Agilent 2100 Bioanalyzer system. For the orthotopically transplanted KCP^mut^ tumors, snap frozen tissue (cohort C versus vehicle) was homogenized, and RNA was isolated with Maxwell® 16 LEV simplyRNA Purification Kit from Promega. Sample purity was evaluated by nanoDrop and RNA quality validated by agarose gel electrophoresis. Transcript levels were quantified with Kallisto (v0.46), using the GENCODE reference transcriptome (mouse version m25 and human version h34). For the human cell line xenograft samples, the human and mouse reference transcriptome were combined, and only human transcripts were kept for downstream analysis. The transcript levels were summed to gene levels and the gene expression levels were normalized between samples with EdgeR (v3.26.8) using trimmed mean of M-values. MAPK activity scores were calculated as described (44), using the normalized log2 counts per million values. For the following genes: SPRY2, SPRY4, ETV4, ETV5, DUSP4, DUSP6, CCND1, EPHA2, and EPHA4.

### Protein lysate preparation and immunoblotting

Cells were plated in complete medium. The morning after, cells were refreshed with medium and drugs of interest. At the desired time points, the cells were washed with cold-PBS and lysed in RIPA buffer supplemented with Halt™ Protease & Phosphatase single-use inhibitors cocktail (100 X) (78442) and Halt™ Protease single-use inhibitors cocktail (100 X) (78430). Protein quantification was performed with the BCA Protein Assay Kit (Pierce). The lysates were then resolved by electrophoresis in Bolt 4 – 12% Bis-Tris Plus Gels (Thermo Fisher Scientific) followed by western blotting as described previously (25). The following antibodies were used: Antibodies against RSK (8408) and phosphorylated RSK (9344 and 8753) were purchased from Cell Signaling Technology (CST). Antibody against alpha-Tubulin (T9026) was purchased from Sigma-Aldrich. Relative pRSK1 levels were quantified by densitometry using ImageJ.

### Statistics

All *in vitro* data are expressed as averages from at least two technical replicates ± SD, unless differently stated, and they have been independently reproduced at least twice with similar results. Significance was determined by ordinary one-way ANOVA test or by one-way ANOVA with Bonferroni’s multiple comparison test or by unpaired, two-tailed t test. Statistical analysis was performed with GraphPad PRISM 8.0 software.

### Study Approval

All animal experiments and care were performed in accordance with the guidelines of institutional committees and European regulations (Directive 2012/63/UE) and approved by the local authorities, Regierung von Oberbayern (ROB-55.2-2532.Vet_02-15-143), the animal experiment committee at the Netherlands Cancer Institute (IVD 1.1.9082), the Dutch Central Authority for Scientific Procedures on Animals (AVD30100202010644), the Universidad Autónoma de Madrid Ethics Committee (CEI 60-1057-A068) and the Comunidad de Madrid (PROEX 335/14).

## Author contributions

The authors confirm contribution to the paper as follows: RB, HA, SM, ML, BS, KJC, ABe and AMS designed the study. *In vitro* cell culture and colony formation assays were performed by AMS, SM and ABo. Apoptosis assay and immunoblotting were done by AMS. *In vitro* data was analyzed by KJC and AMS. KJC, ABe, ML, KG, NW, KND, EK-A, FS, and LH conducted the KCP mouse model experiment. KJC and ABe performed the subcutaneous mouse model experiment. SM, MvdV and NP designed and performed the human xenograft mouse model experiment. LRC conducted the orthotopic mouse model experiment. Human PDX mouse model experiment was done by LRC. Imaging (MRI and PET) was performed by KJC, ABe, KND, NW and KG with the help of PICTUM. KJC analyzed the MRI data. SK analyzed the PET data. RNA isolation and MAPK activity scoring were done by ABe, SM and BT. Histology was performed by ABe. Image acquisition and quantification was performed by ML and ABe. Maintenance of mouse strains and genotyping was done by ABe, KJC, NW, KG, EK-A, DK, KND, YF, FS, LH, MvdV and NP. KJC, ABe, AMS, BS, ML, and SM selected and analyzed the data shown in this manuscript. ML generated the figures with the help of ABe and KJC. KJC, ABe, AMS, ML, and SM wrote the manuscript. The manuscript was edited by DAR, BS, SK, HA, and RB. Supervision was provided by RB and HA. The shared first authors KC, AMS and ABe contributed equally to the experimental work, and their order was assigned considering that KC and AMS also produced preliminary data, and KC coordinated inter-lab meetings.

## Acknowledgments

We thank the Preclinical Imaging Core at TranslaTUM (PICTUM) at Klinikum rechts der Isar der Technischen Universität München for its support, specifically Markus Mittelhäuser and Hannes Rolbieski. We thank the NKI Intervention Unit for technical assistance with the human cell lines xenograft experiment. This work was funded by the American Association for Cancer Research, Lustgarten Foundation and Stand up to Cancer as a Pancreatic Cancer Collective New Therapies Challenge Grant (GRANT NO.SU2C-AACR-PCC-01-18).

