## Supplementary Figures for "Combined SHP2 and ERK inhibition for the treatment of KRAS-driven Pancreatic Ductal Adenocarcinoma"

### Supplemental material

#### Figure S1

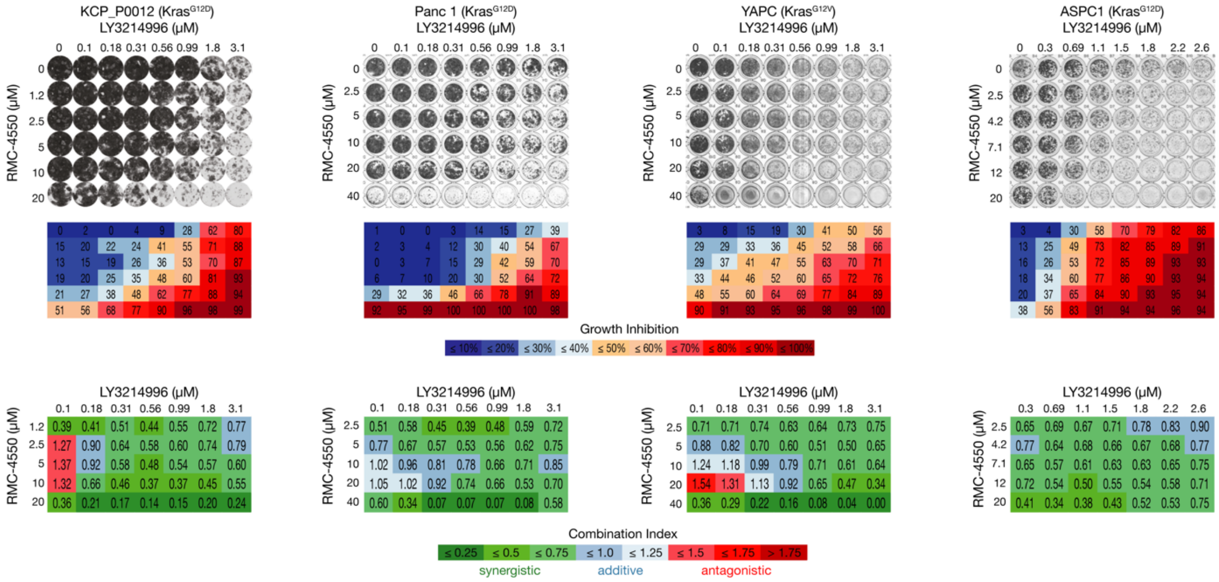


**Supplementary Figure 1. Assessing the treatment response in murine and human *KRAS*-mutant pancreatic cancer cell lines**

Synergistic effects of SHP2i and ERKi administration were evaluated by colony formation assay with murine cancer cell line KPC_P0012 derived from KPC mouse model ($Kras^{G12D}$) of spontaneous tumor formation and human cancer cell lines: Panc 1 ($KRAS^{G12D}$), YAPC ($KRAS^{G12V}$), and ASPC1 ($KRAS^{G12D}$). SHP2i and ERKi were combined at the concentrations indicated. Representative crystal violet staining of cells is shown (top panel). Box-matrices below the plate-scans depict quantification of growth inhibition in relation to control wells (middle panel). Bottom panel: Calculation of the Combination Index (CI) Scores from the growth inhibition values (shown above) via CompuSyn software demonstrating strong synergism between SHP2i and ERKi across a wide range of combinatorial concentrations. CI < 0.75 (shades of green) indicates synergism, CI = 0.75 – 1.25 (shades of blue) indicates additive effects and CI > 1.25 (shades of red) indicates antagonism. Experiments were repeated independently at least three times each, with similar results.

#### Figure S2

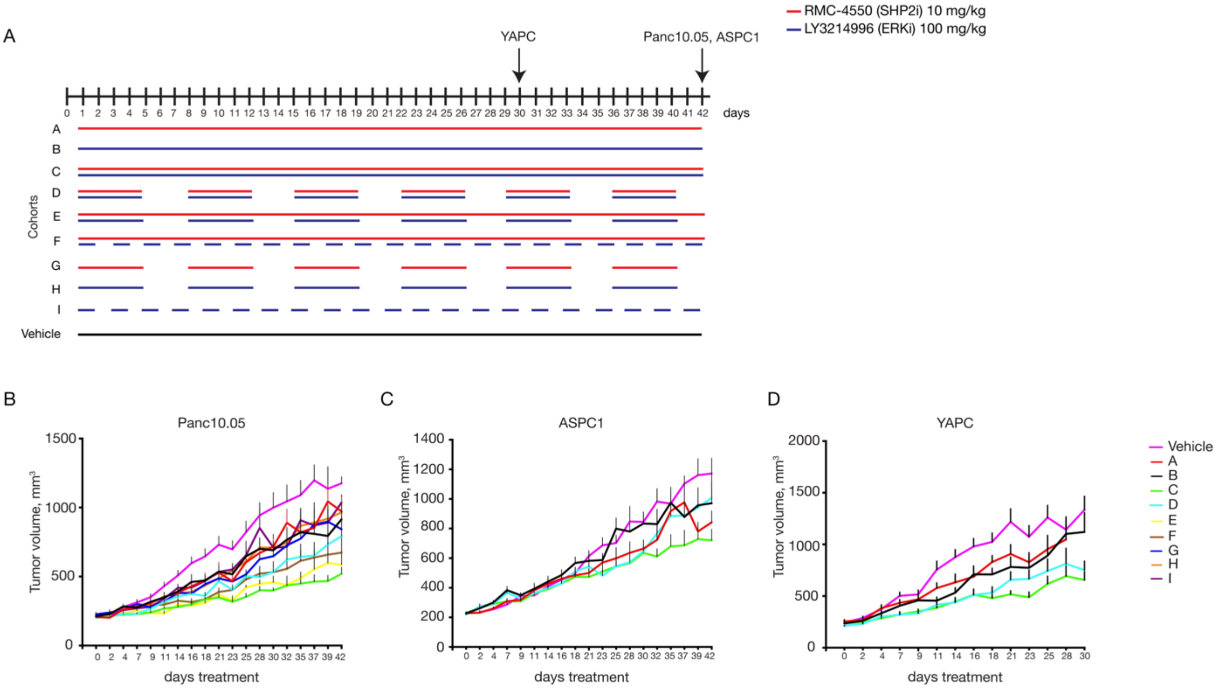


**Supplementary Figure 2. *In vivo* evaluation of the combined administration of RMC-4550 (SHP2i) and LY3214996 (ERKi) on tumor growth in different xenograft models**

**A:** Schematic representation of the treatment schedule applied in MiaPaCa-2, ASPC1, and YAPC xenograft models. Cohort A: Continuous treatment with SHP2i alone daily; Cohort B: Continuous treatment with ERKi alone daily; Cohort C: Continuous treatment with the combination of SHP2i and ERKi daily; Cohort D: Intermittent treatment with the combination of SHP2i and ERKi 5 days on / 2 days off; Cohort E: Semi-continuous treatment schedule with daily dosing of SHP2i and intermittent dosing with ERKi 5 days on / 2 days off. Cohort F: Continuous treatment with SHP2i and on alternate days with ERKi. Cohort G: Intermittent dosing with SHP2i alone 5 days on / 2 days off. Cohort H: Intermittent dosing with ERKi alone 5 days on / 2 days off. Cohort I: Treatment with ERKi alone on alternate days. Control mice were continuously treated with vehicle. For all the xenograft experiments, $5 \times 10^6$ cells were subcutaneously injected into the right flank of NOD scid gamma (NSG) mice, respectively. When tumors reached 200 - 250 $mm^3$, mice were randomly assigned into cohorts and treated by oral gavage with inhibitors or vehicle according to treatment schedule. **B:** Treatment response was assessed through tumor volume change using caliper measurements 3 times/week in different xenograft models: Panc 10.05 ($KRAS^{G12D}$), ASPC1 ($KRAS^{G12D}$), and YAPC ($KRAS^{G12V}$). Results represent mean ± SD.

**Figure S3**

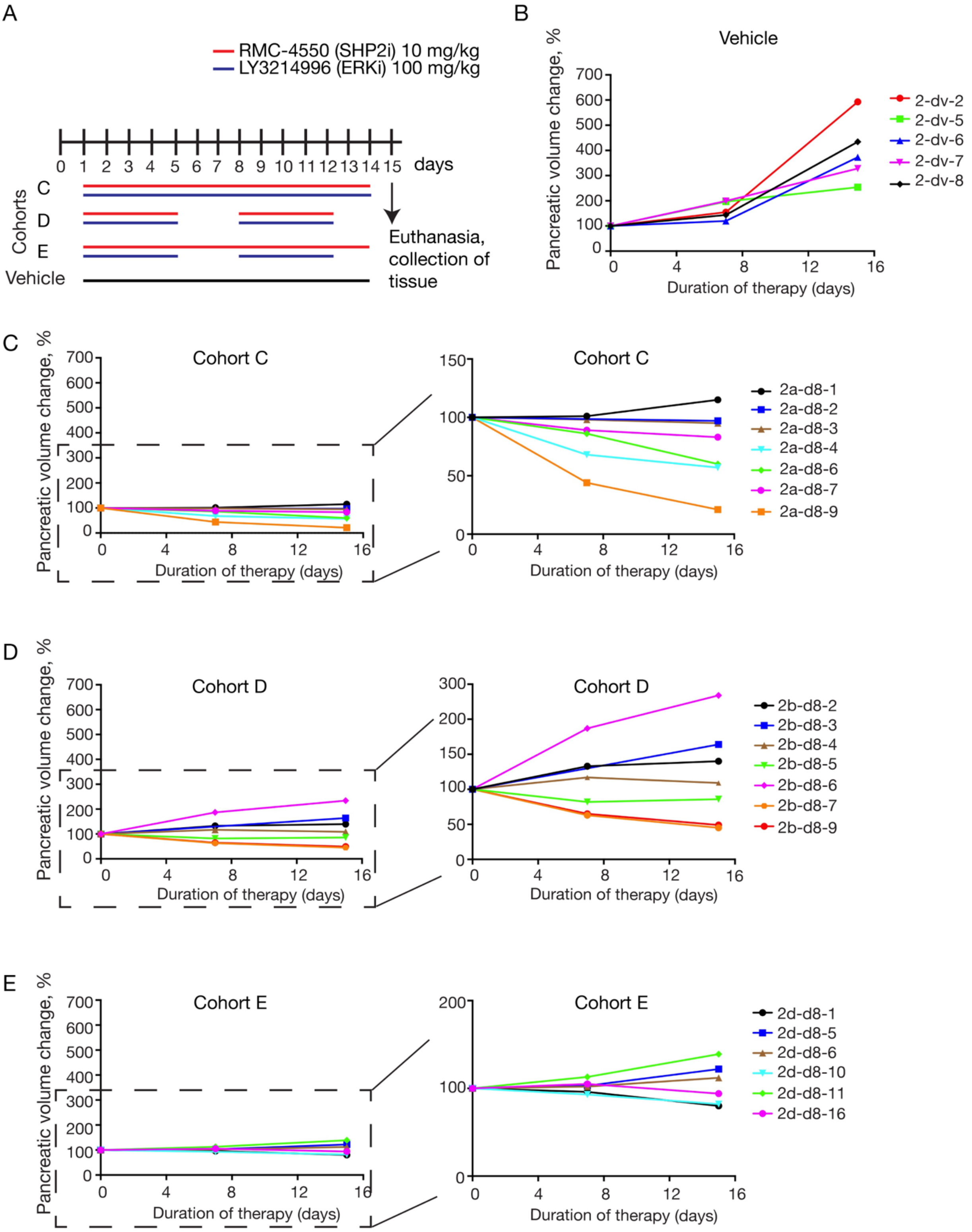
A
RMC-4550 (SHP2i) 10 mg/kg
LY3214996 (ERKi) 100 mg/kg
0 1 2 3 4 5 6 7 8 9 10 11 12 13 14 15 days
Cohorts
C
D
E
Vehicle
Euthanasia, collection of tissue
B
Vehicle
Pancreatic volume change, %
Duration of therapy (days)
2-dv-2
2-dv-5
2-dv-6
2-dv-7
2-dv-8
C
Cohort C
2a-d8-1
2a-d8-2
2a-d8-3
2a-d8-4
2a-d8-6
2a-d8-7
2a-d8-9
D
Cohort D
2b-d8-2
2b-d8-3
2b-d8-4
2b-d8-5
2b-d8-6
2b-d8-7
2b-d8-9
E
Cohort E
2d-d8-1
2d-d8-5
2d-d8-6
2d-d8-10
2d-d8-11
2d-d8-16

**Supplementary Figure 3.** ***In vivo*** **assessment of treatment response in an endogenous murine PDAC model**

**A**: Schematic representation of the treatment schedule applied in an endogenous (KPC) murine model of spontaneous tumor formation. Cohort C: Continuous treatment with the combination of SHP2i and ERKi daily; Cohort D: Intermittent treatment with the combination of SHP2i and ERKi 5 days on / 2 days off; Cohort E: Semi-continuous treatment schedule with daily dosing of SHP2i and intermittent dosing with ERKi 5 days on / 2 days off. Control mice were treated with vehicle for 14 consecutive days. **B - E**: Volume-tracking curves for individual mice over the whole course of therapy. The y axis shows tumor volume change in %. Tumor volume at baseline before commencement of therapy is indicated by 100 %.

**Figure S4**

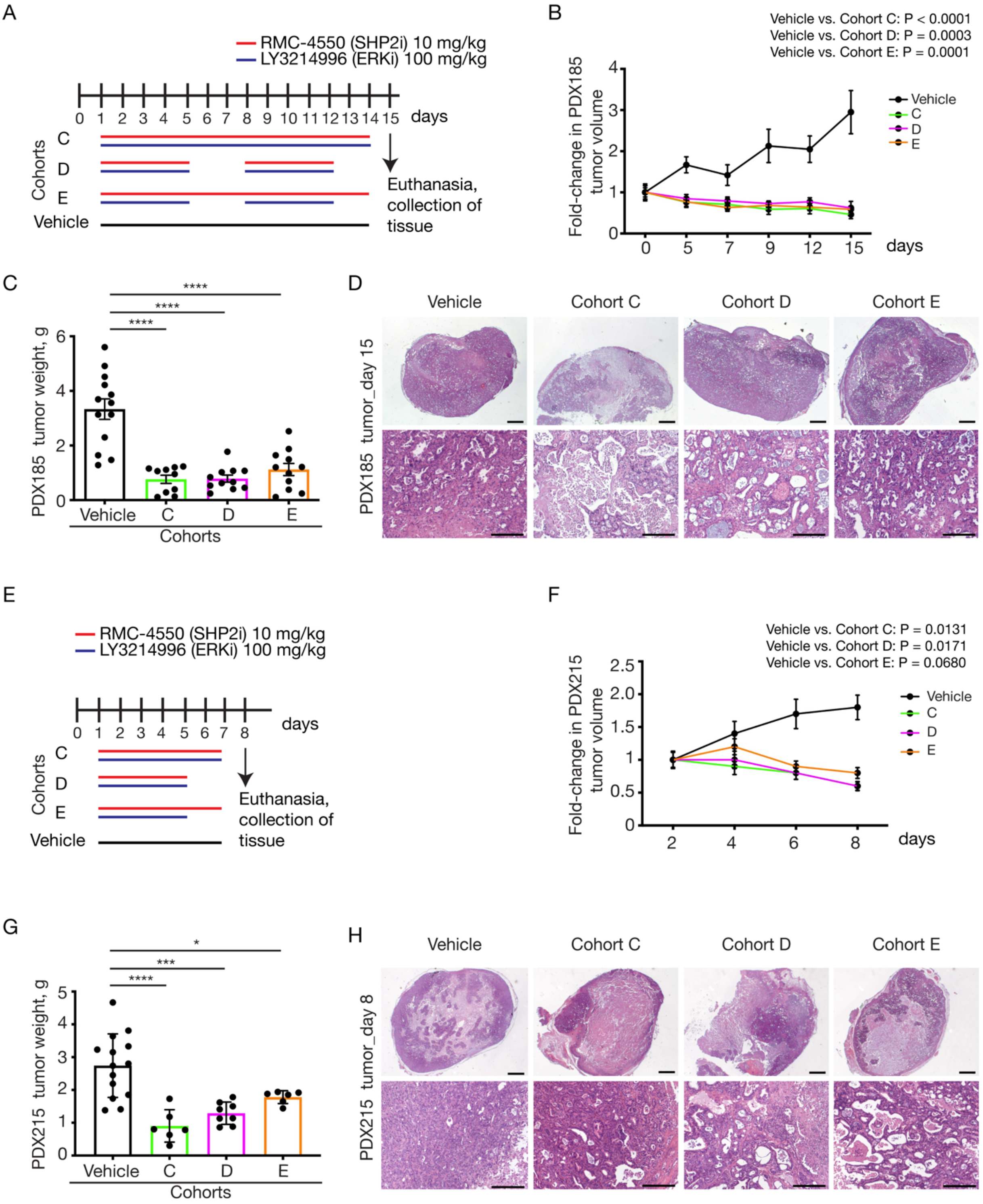
A
RMC-4550 (SHP2i) 10 mg/kg
LY3214996 (ERKi) 100 mg/kg
0 1 2 3 4 5 6 7 8 9 10 11 12 13 14 15 days
Cohorts
C
D
E
Vehicle
Euthanasia, collection of tissue
B
Vehicle vs. Cohort C: P < 0.0001
Vehicle vs. Cohort D: P = 0.0003
Vehicle vs. Cohort E: P = 0.0001
Fold-change in PDX185 tumor volume
Vehicle
C
D
E
days
C
PDX185 tumor weight, g
****
Vehicle
C
D
E
Cohorts
D
Vehicle
Cohort C
Cohort D
Cohort E
PDX185 tumor_day 15
E
RMC-4550 (SHP2i) 10 mg/kg
LY3214996 (ERKi) 100 mg/kg
0 1 2 3 4 5 6 7 8 days
Cohorts
C
D
E
Vehicle
Euthanasia, collection of tissue
F
Vehicle vs. Cohort C: P = 0.0131
Vehicle vs. Cohort D: P = 0.0171
Vehicle vs. Cohort E: P = 0.0680
Fold-change in PDX215 tumor volume
Vehicle
C
D
E
days
G
PDX215 tumor weight, g
*
***
****
Vehicle
C
D
E
Cohorts
H
Vehicle
Cohort C
Cohort D
Cohort E
PDX215 tumor_day 8

**Supplementary Figure 4. Treatment response in Patient-Derived Xenograft (PDX) models**

**A**: Treatment schedule for the PDX185 model. Mice were treated with the combination of RMC-4550 (SHP2i) and LY3214996 (ERKi) once per day via oral gavage for 14 consecutive days (Cohort C) or 5 days on / 2 days off (Cohort D) or with SHP2i continuous and ERKi 5 days on / 2 days off (Cohort E). For the PDX models, tumor pieces of 50 $mm^3$ were subcutaneously implanted into both flanks of NSG mice. When tumors reached 200 - 250 $mm^3$ (approximately 6 - 8 weeks after subcutaneous transplantation), mice were randomly assigned into cohorts and treated by oral gavage with inhibitors or vehicle according to treatment schedule for the indicated times. **B** - **C**: Treatment response of PDX185 was assessed through tumor volume changes using daily caliper measurements (**B**) and tumor weight at endpoint (**C**). Results represent mean ± SD. **** Significance was determined by one-way ANOVA with Bonferroni's multiple comparison test. **D**: Representative H&E-stained sections of vehicle- and combination therapy-treated PDX185 tumors at day 15. Scale bars represent 1000 µm (top) and 200 µm (bottom). **E**: Treatment schedule for the PDX215 model. Mice were treated with the combination of RMC-4550 (SHP2i) and LY3214996 (ERKi) once per day via oral gavage for 7 consecutive days (Cohort C) or 5 days on / 2 days off (Cohort D) or with SHP2i continuous and ERKi 5 days on / 2 days off (Cohort E). **F-G**: Treatment response of PDX215 was assessed through tumor volume changes using daily caliper measurements (**F**) and tumor weight at endpoint (**G**). Results represent mean ± SD. Significance was determined by one-way ANOVA with Bonferroni's multiple comparison test. **H**: Representative H&E-stained sections of vehicle- and combination therapy-treated PDX215 tumors at day 8. Scale bars represent 1000 µm (top) and 200 µm (bottom).

#### Figure S5

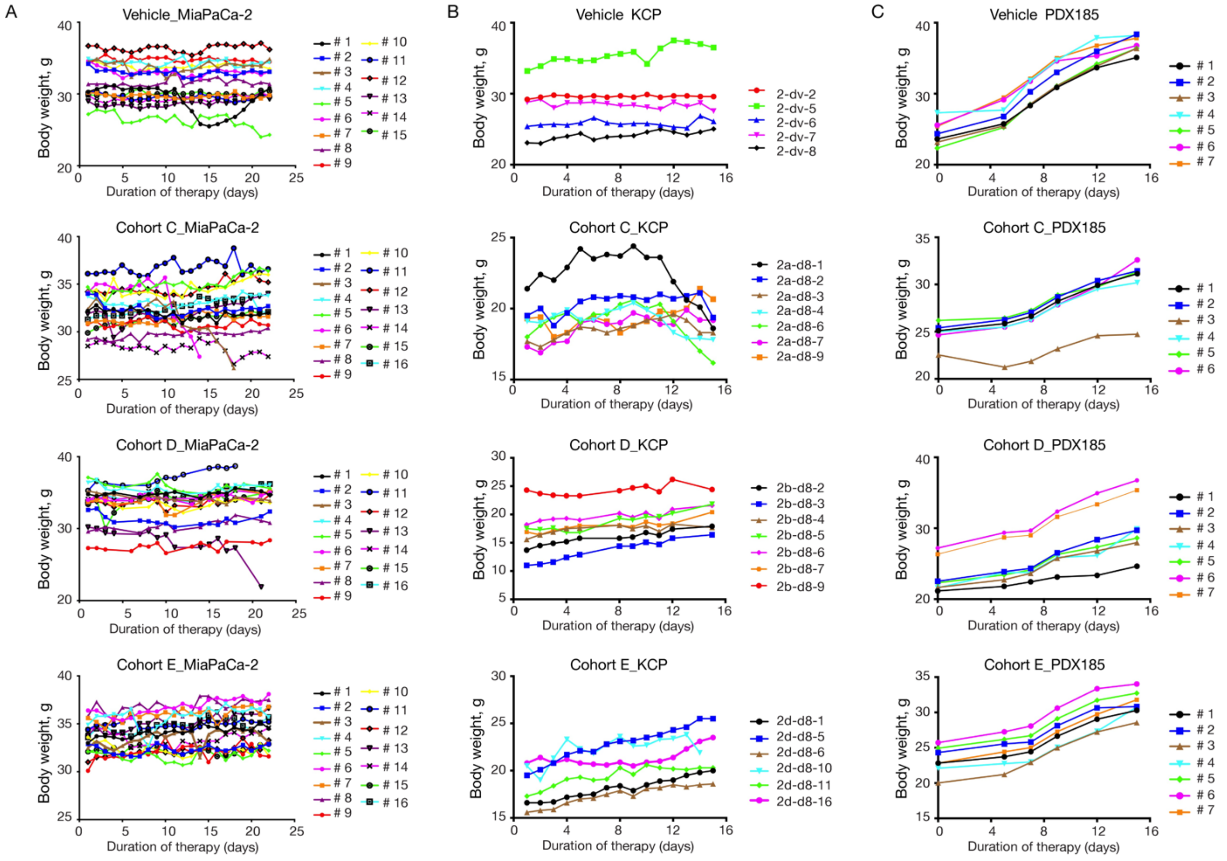


**Supplementary Figure 5. Comparison of body weight–time profiles between the different combination therapy cohorts in the endogenous murine PDAC model, the MiaPaCa-2 xenograft as well as the PDX model**

**A**: Individual body weight–time profile of the vehicle and combination therapy groups in MiaPaCa-2 xenograft bearing mice: Vehicle (n = 15), Cohort-C (n = 16), Cohort-D (n = 16), Cohort-E (n = 16). **B**: Individual body weight–time profile of the vehicle and combination therapy groups in the endogenous (KPC) murine model of spontaneous tumor formation: Vehicle (n = 5), Cohort-C (n = 7), Cohort-D (n = 7), Cohort-E (n = 6). **C**: Individual body weight–time profile of the vehicle and combination therapy groups in PDX185 bearing mice: Vehicle (n = 7), Cohort-C (n = 6), Cohort-D (n = 7), Cohort-E (n = 7).
